# Reproductive competition amplifies purifying selection and reduces genetic load

**DOI:** 10.64898/2026.11.20.768053

**Authors:** Cock van Oosterhout, Jack N. Marsters, Thomas Birley

## Abstract

Deleterious mutations impose genetic load, and purging them through hard selection can carry substantial demographic costs. Competitive and sexual selection can instead remove deleterious variation through differential reproductive success. What remains unclear is when this process is more efficient than equivalent hard selection, and how competition generates this advantage. Here, we use forward-time simulations to express the impact of competition as an effective selection coefficient (*s_e_*) calibrated against hard selection. Competition with multiple rivals strongly amplified purifying selection, whereas repeated contests improved discrimination by averaging stochastic and environmental variation. Strong competition also reduced genetic load relative to hard selection. These effects can shift weakly deleterious mutations from drift-dominated behaviour towards efficient purging. By front-loading purifying selection into reproduction, competitive selection can reduce genetic load at lower demographic cost. This provides a population-genetic mechanism by which genic capture can translate genome-wide genetic quality into more effective purifying selection. This may be particularly important in bottlenecked populations, where demographic recovery can relax purifying selection and allow genomic erosion to persist.

## 1. Introduction

Deleterious mutations impose genetic load by reducing individual fitness and population performance. Purifying selection limits their accumulation, but selection against weakly deleterious mutations becomes less effective in small or bottlenecked populations (Dussex et al. 2023). When purging acts through viability, it can also impose a substantial demographic cost by removing individuals from the population. Sexual and competitive selection can remove deleterious variation through differential reproductive success, potentially at lower demographic cost (Whitlock 2000; Siller 2001; Whitlock and Agrawal 2009; Agrawal and Whitlock 2012). Theory and experiments show that sexual selection can reduce genetic load and increase population fitness (Almbro and Simmons 2014; Lumley et al. 2015; Noël et al. 2019; Cally et al. 2019). Recent genomic evidence links stronger sexual selection to fewer deleterious variants and lower extinction risk under inbreeding (Pointer et al. 2026). However, environmental variation and reproductive skew can weaken this association and allow greater accumulation of deleterious alleles through drift (Tschol et al. 2023). What remains unresolved is how the mode of selection changes the evolutionary fate of deleterious mutations. In particular, we lack a quantitative understanding of when reproductive competition weakens or amplifies purifying selection, which features of competition drive that effect, and how this translates into genetic load. These processes may be particularly important in bottlenecked populations, where genomic erosion can reduce fitness during demographic recovery (Wang et al. 2026a).

The efficacy of sexual and competitive selection depends on how reliably reproductive success reflects underlying genetic fitness. Genic capture allows conditionhdependent traits to reflect realised load caused by deleterious variants, exposing genetic quality to sexual selection (Rowe and Houle 1996; Siller 2001; Whitlock and Agrawal 2009). Under hard selection, fitness acts directly on survival or reproduction, whereas competitive selection allocates a fixed number of reproductive opportunities according to relative performance (Wallace 1975; Reznick 2016; Bell et al. 2021). Relative competition can therefore magnify small fitness differences when competitive performance is closely coupled to reproductive success. Under competitive selection, encounters with multiple rivals assess relative performance more broadly, whereas repeated contests can average stochastic outcomes and transient environmental variation. Multiple and repeated competitive interactions occur naturally in mating, territorial and other contests (Arnold and Wade 1984; Hsu et al. 2006; Hunt et al. 2009; McGlothlin 2010). The same genotype-level fitness effect may therefore have very different evolutionary consequences under different modes of selection.

To compare these modes quantitatively, we separate the assigned selection coefficients (*s*) of variants from their realised population-genetic effects on observed allele-frequency change. In standard models, *s* describes the relative fitness effect of a mutation or genotype. However, allele-frequency change also depends on how fitness differences are translated into survival and reproduction (Charlesworth 2009; Thurman and Barrett 2016). We therefore define an effective selection coefficient, *s_e_*, as the coefficient under hard selection that gives a mutation the same probability of reaching a specified allele frequency as under the focal selection regime. Similar effective coefficients have been used to describe fluctuating selection and population structure (Ohta 1972; Cherry and Wakeley 2003). Comparing *s_e_* with *s* provides a common scale for asking whether a mode of selection weakens or amplifies the evolutionary effect of a given deleterious mutation.

Weakly deleterious mutations around the nearly neutral boundary are particularly sensitive to these effects. When selection is weak relative to drift, small changes in effective strength can substantially alter the evolutionary fate of mutations. A newly arisen deleterious mutation occurs in heterozygotes, so selection initially acts through its heterozygous effect, *hs*. After a bottleneck, increased homozygosity changes the role of dominance. For recessive and partially recessive mutations, much of the deleterious effect is masked in heterozygotes but fully expressed in homozygotes. Under standard hard selection, López-Cortegano and Charlesworth (2026) show that purging requires 2*N*e|*s*| to be substantially greater than one and *h* < 1/3. Below this dominance threshold, increased homozygosity exposes recessive deleterious effects in homozygotes strongly enough to increase the net removal of mutant alleles. These results show that the efficacy of purging depends not only on the intrinsic effect of a mutation, but also on how that effect is expressed and translated into selection across heterozygous and homozygous genotypes.

Here, we use forward-time simulations to test how different modes of selection alter the fate of deleterious mutations and the purging of genetic load. We compare hard selection with competitive selection while varying the number of distinct rivals and the number of repeated contests. We also compare this with population-wide rank selection, representing maximum competitive assessment. We use *s_e_* to place their effects on a common hard-selection scale. We also test how transient environmental variation affects competitive discrimination and whether repeated assessment recovers persistent differences in genetic fitness. We predict that competition will amplify purifying selection, with amplification increasing as individuals encounter more rivals and repeated contests reduce noise. We then test whether these changes in effective selection translate into differences in the site-frequency spectrum and genetic load.

## 2. Results

### 2.1 Competition can weaken or amplify effective selection

Under absolute selection, the probability that a new deleterious mutation reached *q* = 0.05 declined as |*s*| increased. This relationship provided the calibration used to estimate *s*_e_ for the competitive treatments. By definition, hard selection represents the reference *s*_e_ = *s*, see Methods. Competitive selection produced departures from this reference in both directions. With one rival and one contest round (*D* = 1, *R* = 1), |*s*_e_| was only about 0.4–0.6 times the assigned |*s*|. Nine rounds against the same rival (*D* = 1, *R* = 9) increased |*s*_e_| to approximately |*s*| (Figure 1A). Increasing the number of distinct rivals had a much stronger effect. With one round per rival, *D*=5 produced |*s*_e_|/|*s*| ≈ 1.4–1.8 and *D* = 10 produced ≈ 2.1–2.3. Combining multiple rivals with repeated contests produced the strongest amplification; at *D* = 10 and *R* = 9, |*s*_e_| was approximately five to six times the assigned genotype-level effect (Fig. 1a; Fig. S4). These differences in effective selection translated into marked differences in genetic load: weak competition (*D* = 1, *R* = 1) produced the greatest accumulation, whereas increasing competitive intensity progressively reduced load, with *D* = 10, *R* = 9 maintaining the lowest levels (Fig. 1b). The allele-frequency distribution showed the same pattern: weak competition allowed more deleterious mutations to reach high frequencies and fixation, whereas strong competition concentrated them at very low frequencies and largely prevented progression towards fixation (Fig. 1c–f).

**Figure 1.**
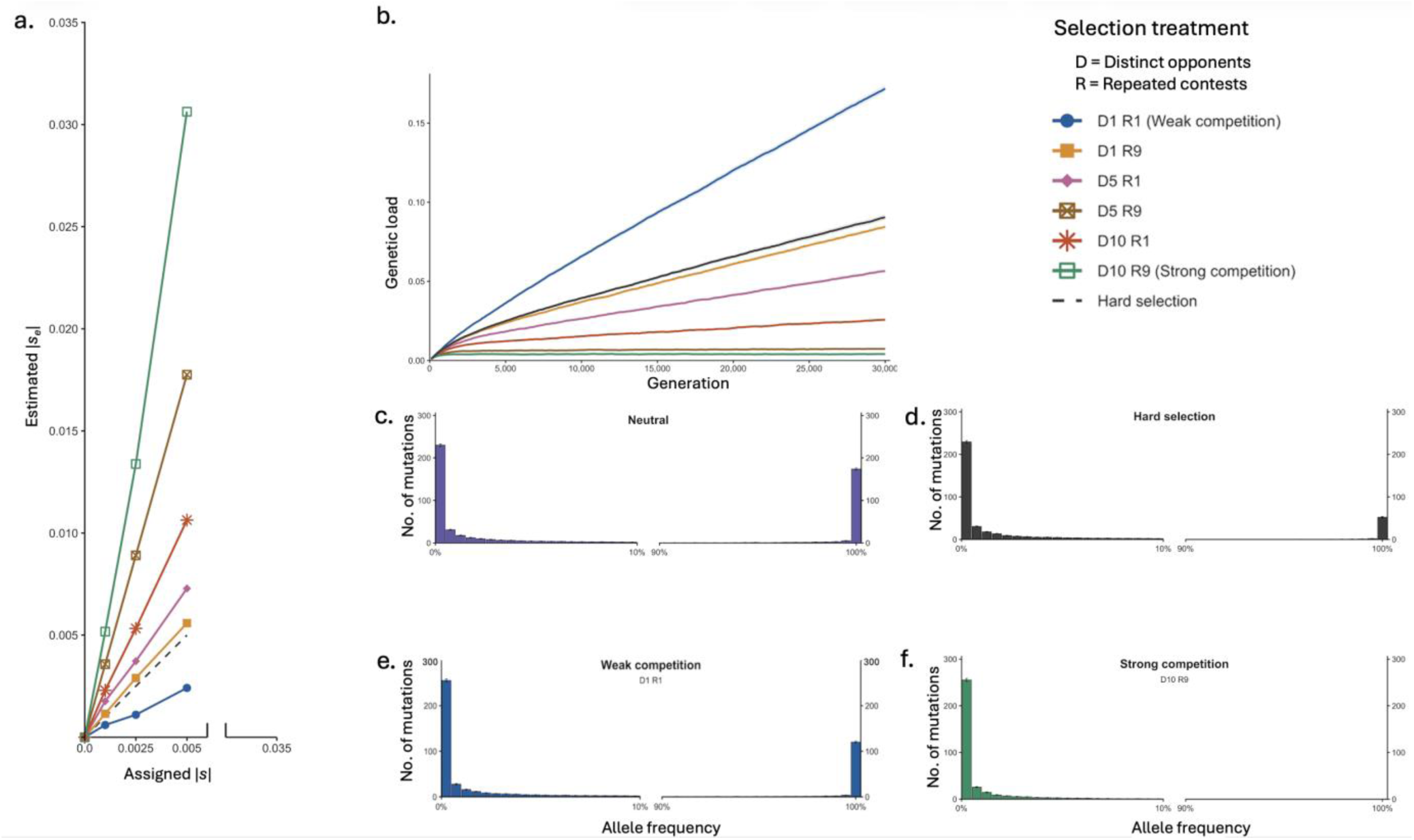
Competitive selection alters the effective strength of selection and the accumulation of deleterious variation. a, Assigned genotype-level selection coefficient, ∣*s*∣, versus the estimated effective selection coefficient, ∣*s_e_*∣, under six competitive-selection regimes differing in the number of distinct opponents (*D* = 1, 5, 10) and repeated contests per opponent (*R* = 1, 9). The dashed line denotes the hard-selection expectation ∣*se*∣=∣*s*∣; values below and above this line indicate attenuation and amplification of selection, respectively. b, Mean genetic load through 30,000 generations for deleterious mutations with *s* = −0.001 and ℎ = 0.5 under hard selection and the six competitive-selection regimes. Lines show means across 100 replicate populations. c–f, Allele-frequency distributions at generation 30,000 under c, neutrality (*s* = 0); d, hard selection (*s* = −0.001); e, weak competition (*D* = 1, *R* = 1); and f, strong competition (*D* = 10, *R* = 9). Bars show the mean number of mutations across replicate populations. Only the 0–10% and 90–100% frequency ranges are shown; the terminal class (*q* ≥ 0.995) includes mutations at or near fixation.

### 2.2 Increasing the number of distinct rivals strengthens effective selection

To examine this effect of multiple rivals in further detail, we set the number of repeat contests to one (*R*=1) whilst holding the environmental variation constant (*σ*_E_ = 0.25). We examined increasing the number of distinct rivals from *D* = 1 to 100. Effective selection increased progressively with *D*; competition with one or two rivals weakened selection relative to hard selection, whereas five or more rivals amplified it (Fig. 2A). The amplification increased to approximately fourfold at *D* = 100, while population-wide ranked selection produced an approximately 5.6-fold increase in |*s*_e_|/|*s*|. Thus, broader sampling of relative fitness across independent rivals progressively strengthened effective selection and approached the effect of population-wide ranking.

**Figure 2.**
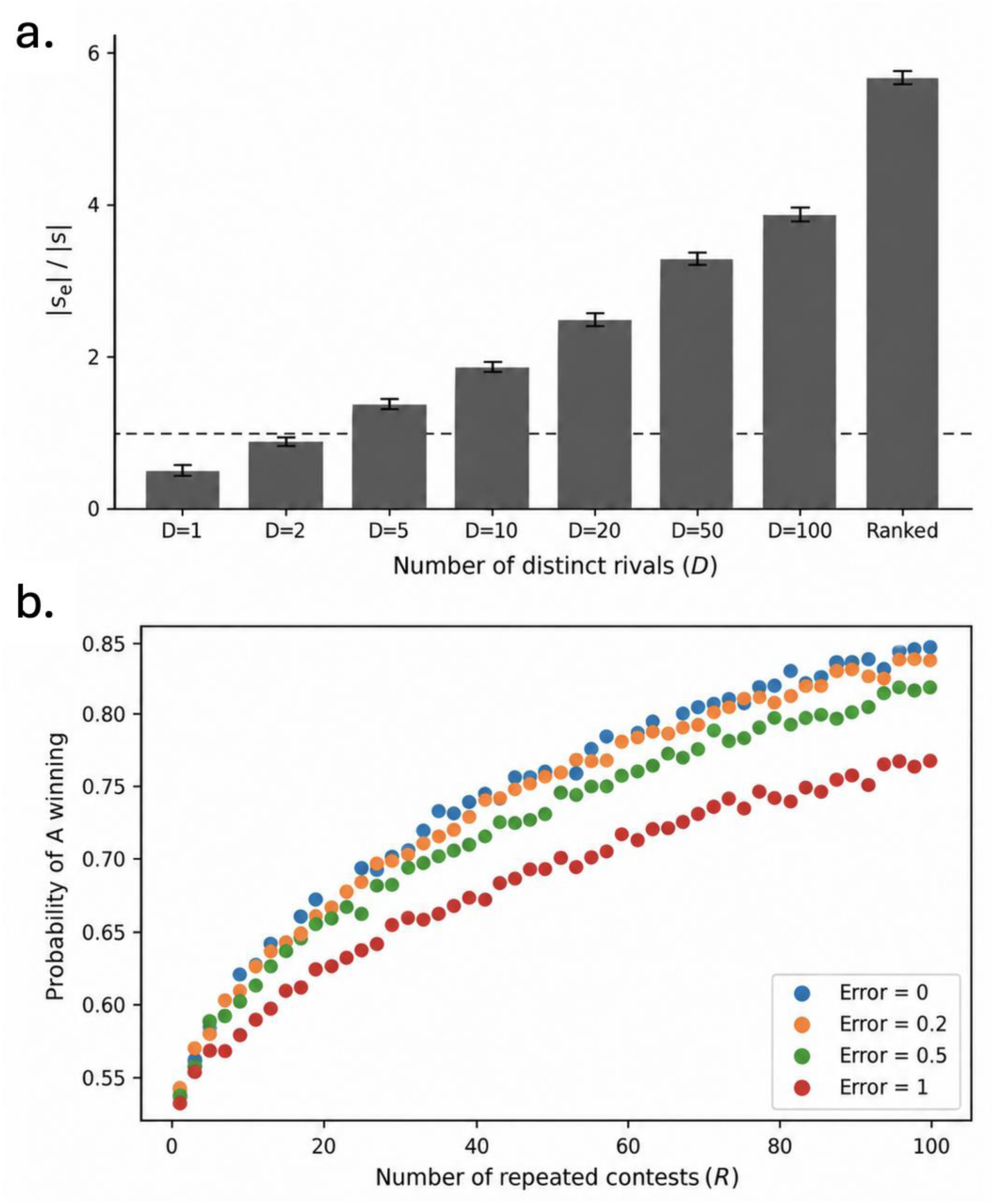
Competitive structure shapes how fitness differences translate into effective selection. **(a)** Effect of the number of distinct rivals on the relative strength of competitive selection under environmental variation. Bars show the ratio of the effective selection coefficient to the assigned selection coefficient (|*s_e_*|/|*s*|) for Bernoulli competition with different numbers of distinct rivals, as well as for ranked selection. Simulations used *R* = 1, *σ*_E_ = 0.25, assigned s = -0.0025 and ℎ = 0.5. Bars show the mean |*s_e_*|/|*s*| and error bars show bootstrap 95% confidence intervals. The dashed horizontal line indicates |*s_e_*|/|*s*| = 1. (b) Repeated contests reduce the effect of transient environmental variation. Shown is the probability that the fitter individual A wins the majority of a contest series as the number of contests increases, for Δ*L* = 0.2 and four levels of transient environmental noise. Repeated assessment progressively improved discrimination between competitors, even under substantial environmental noise.

### 2.3 Scaling and robustness of the effective-selection calibration

Within each competitive regime, |*sₑ*| increased approximately linearly with |*s*| over the range 0.001–0.005, and the ratio |*sₑ*|/|*s*| was broadly stable (Fig. S4). Thus, over this range, the competitive regime acted approximately as a multiplier of genotype-level fitness effects. Changing *K* altered *Nₑ*, but had little effect on the probability that neutral mutations reached *q* = 0.05 (Fig. S1), suggesting that differences in drift were unlikely to explain the much larger changes in *sₑ*. Additional sensitivity analyses across candidate population sizes (*N_c_* = 100–25,000) showed that selection amplification was broadly robust for *N_c_* ≥ 500, particularly after accounting for differences in neutral threshold-crossing probabilities between selection regimes (Fig. S7; Tables S1, S2). Estimates were less reliable in the smallest, drift-dominated populations (*N_c_* = 100 and 250). Likewise, random assignment of the same skewed reproductive weights did not strengthen selection (Fig. S8), showing that reproductive skew alone was insufficient. Amplification instead depended on coupling reproductive success to genotype-dependent competitive performance. Competition treatments and their matched hard-selection references showed broadly similar *q_max_* distributions across the full range of allele frequencies (Fig. S2), and this correspondence was retained across dominance coefficients from *h* = 0 to 0.5 (Fig. S3). Additional sensitivity analyses showed that weak competition generally attenuated purifying selection, whereas stronger competition amplified it across dominance coefficients. Expressing selection in terms of heterozygous fitness effects (*hs*) produced the same qualitative pattern, although estimates were less precise for strongly recessive mutations (Figs. S5, S6). The contrast between weak and strong competition was also retained at higher allele-frequency thresholds and at fixation (Fig. S11). Selection amplification was also broadly maintained across 100-fold variation in mutation rate and the recombination parameter, *ρ* (Tables S3, S4).

### 2.4 Repeated contests reduce the effect of environmental noise

In the pairwise analysis, the fitter individual became increasingly likely to win the contest series as the number of contests increased. The simulated increase closely followed the analytical binomial majority-vote prediction (Fig. S9). At *ΔL* = 0.2, its probability of winning rose from approximately 0.55 in a single contest to 0.84 after 99 contests without environmental variation. Adding transient environmental variation reduced discrimination at any given number of contests, but repeated contests progressively recovered the persistent underlying fitness difference (Figure 2B; Figs. S9, S10). Thus, repeated assessment averages stochastic contest outcomes and, when present, short-term environmental variation, allowing persistent fitness differences to determine competitive success more reliably.

## 3. Discussion

The efficacy of purifying selection depends strongly on how selection is realised. For the same genotype-level fitness effect, competitive selection could either weaken selection or amplify its effect. Consequently, at the same assigned *s*, stronger competition reduced genetic load relative to hard selection. We quantify this efficiency of selection to purge deleterious mutations using an effective selection coefficient (*sₑ*): the coefficient under hard selection that gives a new mutation the same probability of reaching *q* = 0.05. Related effective coefficients have previously been used to summarise fluctuating selection and population structure (Ohta 1972; Cherry and Wakeley 2003; Marrec 2026). We found that *sₑ* scaled approximately linearly with *s* across the range examined. This is especially important for weakly deleterious mutations around the nearly neutral boundary, whose fate can be sensitive to relatively small changes in the efficacy of selection (Ohta 1973; de Jong et al. 2024). The evolutionary effect of a mutation therefore depends on both its intrinsic fitness effect and how strongly that effect is exposed during condition-dependent courtship and mate choice. These processes determine how effectively genome-wide genetic quality is captured by selection and ultimately purged from the population.

These results add a population-genetic dimension to the difference between absolute (hard) and competitive (soft) selection. Under soft selection, relative competition determines which individuals reproduce rather than how many, so purging can occur at lower demographic cost than under equivalent hard selection (Wallace 1975; Reznick 2016; Bell et al. 2021). The demographic cost of hard selection can also be buffered by reproductive compensation, where losses of lower-fitness individuals are offset by excess reproduction or density-dependent recruitment (Lesecque et al. 2012; Agrawal and Whitlock 2012; Charlesworth 2013). We show that competitive selection can additionally amplify the evolutionary effect of a given *s*, combining more effective purging with lower demographic cost. When competitive selection acts through mating success, purifying selection can also be front-loaded earlier in the life cycle (Whitlock 2000; Siller 2001; Whitlock and Agrawal 2009). Lower-fitness individuals can lose reproductive opportunities before substantial investment in offspring, while successful competitors compensate reproductively, as expected under soft selection (Bell et al. 2021).

The converse may occur when demographic recovery relaxes selection. Higher juvenile survival and reduced intraspecific competition may weaken purifying selection, as proposed to explain the increase in genetic and realised load in the demographically recovered Seychelles magpie-robin (Cavill et al. 2024). The same mismatch between demographic and genomic recovery has been observed in the whooping crane and Mauritius kestrel, where population recovery did not halt genomic erosion (Fontsere et al. 2025; Wang et al. 2026a). Demographic recovery may therefore fail to restore genomic health when it relaxes the selective processes that purge deleterious variation. By contrast, populations in which purifying selection remains strong during reproductive competition and earlier reproductive stages may maintain lower genetic load and higher viability (van Oosterhout et al. 2022).

Repeated contests and encounters with different rivals affected selection in different ways. Repeated assessment reduced stochastic and environmental noise, whereas multiple rivals sampled relative performance more broadly and produced a much larger increase in *sₑ*. This provides a mechanism linking sexual selection to good-genes and indicator models. Under the genic capture hypothesis, condition-dependent traits integrate genetic effects across many loci, allowing genome-wide deleterious variation to influence reproductive phenotype and thereby become exposed to sexual selection (Rowe and Houle 1996; Siller 2001; Whitlock and Agrawal 2009). Experimental results vary among systems, with sexual selection sometimes reducing mutation load or increasing population fitness and sometimes showing little or opposite effects (McGuigan et al. 2011; Almbro and Simmons 2014; Lumley et al. 2015; Arbuthnott and Rundle 2012; Cally et al. 2019). This heterogeneity may reflect differences in experimental scale and duration, as well as how closely reproductive success tracks genetic fitness.

Recent genomic studies provide clearer support for purging by sexual selection. Comparative studies in birds have shown that stronger sexual selection is associated with more efficient purifying selection. Across 150 species, Wanders et al. (2023) found a lower genome-wide nonsynonymous-to-synonymous polymorphism ratio (*P*_N_/*P*_S_) in polygamous than monogamous species, independent of neutral heterozygosity. Experimental evolution provides more direct evidence. In *Tribolium castaneum*, populations evolving under stronger sexual selection for 156 generations carried fewer deleterious missense and nonsense variants than populations under weak sexual selection, and mutation load predicted subsequent extinction risk under inbreeding (Pointer et al. 2026). This shows that sexual selection can modify genome-wide deleterious variation over evolutionary time, with consequences for population persistence. In a study on wild black grouse, genetic load predicted lower male mating success, with the association acting through lek attendance rather than ornamental traits (Chen et al. 2025). Lek attendance integrates short-term condition with longer-term reproductive effort and is repeatedly exposed to sexual selection across breeding seasons. The authors therefore interpret this relationship in terms of genic capture, whereby genome-wide deleterious variation is expressed through condition-dependent reproductive behaviour. This fits our finding that repeated competitive assessment can resolve small differences in genetic fitness.

Changes in the effective strength of selection are especially consequential for mutations near the nearly neutral boundary. In bottlenecked populations, recent theory under conventional hard selection shows that the outcome of purging depends strongly on *N*_e_, *s* and *h* (López-Cortegano and Charlesworth 2026). Our results extend this framework by showing that the mode of selection can itself alter the effective strength of selection acting on the same underlying mutation. If ecological selection changes the mapping from *s* to *sₑ*, a mutation close to this boundary can move from being dominated by drift to efficient purifying selection without any change in its genotype-level effect (Ohta 1973; de Jong et al. 2024). Conversely, with less competition |*sₑ*| is reduced, making the mutation behave more neutrally. A newly arisen deleterious allele is initially rare and therefore occurs predominantly in heterozygotes, so selection initially acts through its heterozygous effect, *hs*, rather than the homozygous effect *s*. This also explains an apparent factor-of-two difference among formulations of the nearly neutral boundary. The boundary is commonly written as |*s*| ≈ 1/(2*Nₑ*), whereas formulations based on selection against a rare heterozygous allele use |*hs*| ≈ 1/(4*Nₑ*) (Ohta 1973; Charlesworth 2009). For codominant mutations (*h* = 0.5), as used in our calibration of *s* to *sₑ*, these are equivalent.

The genotype-level fitness effect of a molecular change can itself depend on where, when, and how often it is expressed. Highly expressed genes generally evolve more slowly than weakly expressed genes (Pál et al. 2001), and experimental manipulation shows that expression level can alter the fitness effects of coding mutations (Wu et al. 2022). Furthermore, population-genomic analyses infer stronger purifying selection in highly expressed genes (Trucchi et al. 2025). Repeated expression can magnify weak deleterious effects: if the same effect *s* is expressed independently across *r* episodes, cumulativerelative fitness is (1 + *s*)*^r^*, giving a cumulative effect *sᵣ* ≈ *rs*, when |*s*| is small. Thus, competitive selection acts at a different stage, with repeated assessment and exposure to multiple rivals increasing how efficiently a given genotype-level effect is translated into reproductive success, and hence, into *s_e_*. These processes can strengthen selection in two distinct ways: repeated expression can increase the cumulative genotype-level fitness effect of a mutation (*s* → *sᵣ*), whereas competitive assessment can increase how efficiently that effect is translated into allele-frequency change (*s* or *sᵣ* → *sₑ*).

Distinguishing *s* from *s_e_* is important when translating genomic variant-effect predictions into fitness effects and their evolutionary consequences. Variant-effect annotations and deleteriousness scores such as SnpEff, SIFT and CADD provide proxies for the functional effects of mutations (Bertorelle et al. 2022), but these effects must first be translated onto a genotype-level fitness scale, *s*. Recent work on the Mauritius kestrel, for example, combined SnpEff and CADD predictions with simulation-anchored class-specific fitness effects to approximate the mapping from predicted deleteriousness onto a fitness scale (Wang et al. 2026a). Our results show that predicting mutation fate requires a second translation from *s* to *sₑ* because selection mode determines whether a given fitness effect is efficiently purged or persists and continues to contribute to genetic load. To assess how efficiently a mutation is actually selected against, it is informative to compare predicted functional effects with the strength of selection inferred over evolutionary time. Phylogenetic constraint metrics such as GERP reflect the long-term evolutionary consequences of selection and may therefore be more closely aligned with *sₑ* than with the intrinsic genotype-level effect *s*. Variants showing stronger phylogenetic constraint than expected from their predicted functional consequence may represent cases in which |*s_e_*| exceeds that expected from |*s*|. Such variants should therefore be enriched in genes whose fitness effects are repeatedly exposed to strong competitive or sexual selection.

Our results make a clear prediction: weakly deleterious mutations should be purged more efficiently where reproductive competition repeatedly exposes differences in genetic quality. Comparative genomic analyses have already shown that genetic load can be assessed across many species (Wang et al. 2026b). This framework could now be extended by combining whole-genome resequencing with long-term individual fitness data and measures of sexual-selection intensity. Such datasets allow genic capture to be tested directly at the genomic level, linking individual realised load to competitive performance, mate choice and reproductive success. This has direct implications for threatened species and their management. In *ex situ* populations, managed breeding can constrain mate choice and reproductive competition, potentially weakening an important route of purifying selection (van Oosterhout et al. 2026), although genomics-informed pairing may partly compensate for this (Speak et al. 2024). Mating system and competitive behaviour are part of a species’ reproductive ecology, and their disruption can alter how effectively deleterious variation is purged, with consequences for population viability.

## 4. Materials and methods

### 4.1 Forward-time simulations

We used forward-time individual-based simulations in SLiM v4.3.0 under the non-Wright–Fisher framework (Haller and Messer 2023). Simulations contained diploid individuals carrying 1,000 freely recombining loci. The per-locus mutation rate was *μ* = 1 × 10^−5^ per generation. Mutations were neutral or deleterious, with assigned selection coefficients spanning the nearly neutral range and stronger effects. Competition treatments were run with *s* = 0, −0.001, −0.0025 and −0.005, while a broader set of hard-selection coefficients was used to construct the calibration curve. Genetic fitness combined multiplicatively across loci.

For the main analyses, mutations were codominant (*h* = 0.5). This additive reference avoids dominance-dependent non-linearity: each deleterious allele contributes equally to genotype-level fitness, irrespective of whether it occurs in a heterozygous or homozygous genotype. Unless stated otherwise, 100 independent simulation replicates were run for each parameter combination. Additional sensitivity analyses varied dominance (*h* = 0, 0.01, 0.05, 0.1, 0.2, 0.5), candidate population size (*N_c_* = 100, 250, 500, 1,000, 2,500, 5,000, 25,000), mutation rate (*µ* = 10^-6^, 10^-5^, 10^-4^) and recombination parameter (*ρ* = 0.005, 0.05, 0.5). Effective selection was recalibrated against the corresponding hard-selection reference for each treatment (Supplementary Information, Sections S1.4–S1.8).

### 4.2 Absolute (hard) selection

Under absolute selection, the probability that an individual contributed to the next generation depends on its own genotype-level fitness. In our simulations, density regulation acted separately from genotype, so increasing the number of offspring exposed to selection did not itself change the relative fitness disadvantage assigned to a mutation. These simulations provided the hard-selection reference used to calibrate *s_e_* (Section 4.7).

Because differences in effective population size (*N_e_*) alter the strength of drift and hence the probability that a new mutation reaches a given frequency, we aimed to keep *N_e_*approximately comparable between the absolute-selection reference and competitive treatments. In the main simulations, carrying capacity was *K* = 1000, corresponding to an estimated *N_e_* of approximately 1,111 under hard selection and 987 under competitive selection (Table S2). To determine whether this difference in *N_e_*, and hence genetic drift, could explain differences in mutation trajectories, we performed separate calibration simulations in which *K* was varied under hard selection while fecundity was adjusted to maintain an expected offspring cohort of approximately 5000. Across these *K* values, the probability that neutral mutations reached *q* = 0.05 remained similar despite the resulting variation in *N_e_* (Fig. S1). Thus, the much larger differences in threshold-crossing probability between hard and competitive selection are unlikely to be caused by the modest residual difference in *Nₑ* and instead reflect differences in the efficacy of selection.

### 4.3 Competitive selection

Competitive-selection simulations generated a cohort of 5000 candidate offspring each generation. Genotype-level fitness affected reproductive success through contests rather than through an absolute viability threshold. The competition structure was controlled by two parameters: *D*, the number of distinct rivals encountered by each individual, and *R*, the number of repeated contest rounds against each rival. We analysed nine combinations *D* = 1, 5 or 10, and *R* = 1, 3 or 9. Distinct opponents were assigned using a balanced round-robin procedure.

For competitors *i* and *j* with genotype-level fitnesses *w_i_* and *w_j_*, the probability that individual *i* won a single round was

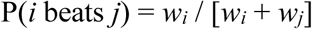

Each pair competed for *R* independent rounds, with the majority winner recorded for that pair. An individual’s competition score was the number of its *D* rivals that it defeated. Individuals were ranked by this score, with ties resolved randomly.

Reproductive opportunity was then allocated using a strongly right-skewed Gamma distribution with shape *α* = 0.245 and scale *θ* = 4.085 (mean ≈ 1; CV ≈ 2.02). Gamma draws were ordered from largest to smallest and assigned by competitive rank, so that higher-ranked individuals received greater expected reproductive success. This parameterisation approximates the strong reproductive skew observed in highly polygynous mammals, including elephant seals, bighorn sheep and Soay sheep (Supplementary Information Table S5). Reproductive weights were normalised to sum to one and used as parental sampling probabilities. Two parents were sampled for each offspring, with self-fertilisation prevented. To distinguish the effect of reproductive skew itself from its coupling to competitive performance, we also ran a control in which reproductive success was drawn from the same Gamma distribution, but the resulting weights were assigned randomly among individuals. This control used *s* = −0.0025, ℎ = 0.5, *D* = 10 and *R* = 9, all other simulation and mutation-tracking procedures unchanged. We compared this treatment with the corresponding competition-linked treatment and a neutral treatment (*s* = 0; Fig. S8).

To examine how the number of distinct rivals affected effective selection more broadly, we ran an additional series of simulations varying *D* (*D =* 1, 2, 5, 10, 20, 50, and 100), with a single contest per rival (*R* = 1) and transient environmental variation fixed at σ*_E_* = 0.25. Before each contest, an independent environmental deviation, ε ∼ *N*(0, σ*_E_*²), was added to each individual’s realised load. Competitive fitness was recalculated from the resulting perceived load, with environmental deviations redrawn independently for each individual and contest. Thus, environmental variation represented transient variation in competitive performance rather than a persistent individual effect. Competitive scores and reproductive success were then determined as described above. We additionally included population-wide rank selection, in which candidate offspring were ranked directly by genotype-level fitness before the same reproductive weights were assigned, representing population-wide assessment of relative fitness.

### 4.4 Contest-level discrimination and environmental noise

To isolate how repeated assessment affects discrimination between two competitors, we ran a separate pairwise analysis using the same contest rule as above. For individuals A and B, fitness was expressed as *w* = exp(−*L*), so the probability of A winning depended only on the difference in realised load, Δ*L* = *L*_B_ − *L*_A_. We focused on a fixed difference of Δ*L* = 0.2, and varied the number of contests in a series from *m* = 1, 3, 5, …, 99. The individual winning the majority of contests was recorded as the overall winner.

To model transient environmental variation in competitive performance, an independent error was added to each individual’s realised load before every contest: *L′* = *L* + ε, with ε ∼ *N*(0, σ*_E_*²). The error was redrawn for every individual and every contest, so it represented short-term variation in condition or performance rather than a persistent environmental effect. We analysed σ*_E_* = 0, 0.2, 0.5 and 1, with 10,000 contest series simulated for each value of *m* and σ*_E_*.

### 4.5 Tracking newly arisen mutations

New mutations were registered from the complete offspring cohort before hard or competitive selection acted, so entry into the analysis was independent of subsequent survival or reproductive success. Mutations arising during generations 10,001–15,000 were tracked. With 5000 diploid offspring, 1000 loci and *μ* = 10^−5^, approximately 100 new mutations were expected per generation, or about 500,000 during the 5000-generation sampling interval in each replicate.

For each mutation we recorded its generation of origin and whether it subsequently crossed allele-frequency thresholds *q* = 0.05, 0.10, 0.20 or 0.50, reached fixation (*q* = 1), was lost, or remained segregating when simulations ended at generation 30,000. A threshold was counted only on its first upward crossing. Mutations still segregating at the end of the simulation were treated as right-censored rather than lost. We also recorded the site-frequency spectrum (SFS) at the end of the simulation. Mutation frequencies were grouped into 0.005-wide bins, with the terminal *q* ≥ 0.995 class including mutations at or near fixation (Fig. 1c-f). Mutation age was calculated from the generation of origin for mutations present at the final sampling point.

### 4.6 Quantification of genetic load

Genetic load was quantified using the individual-based estimator of Bertorelle et al. (2022). For each individual, realised load was calculated as

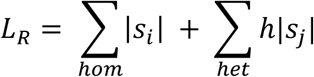

where *s_i_* and *s_j_* are the selection coefficients of deleterious mutations occurring in homozygous and heterozygous genotypes, respectively. In the main simulations, deleterious mutations analysed were codominant (*h* = 0.5); consequently, realised load was equivalent to the individual genetic-load estimator of Bertorelle et al. (2022). Genetic load was calculated for each individual and averaged across individuals within each population. Treatment means were then calculated across 100 independent simulation replicates at each generation. For the long-term analysis (Fig. 1b), we tracked genetic load over 30,000 generations using deleterious mutations with an assigned selection coefficient of *s* = -0.001.

### 4.7 Estimating the effective selection coefficient (*s_e_*)

For each assigned selection coefficient, *s*, we calculated the probability that a newly arisen mutation reached *q* = 0.05 before loss. We then used hard selection as the reference model to determine which hard-selection coefficient produced the same probability. To do this, we fitted a calibration curve relating assigned |*s*| under hard selection to the probability of reaching *q* = 0.05. For each competitive treatment, its observed probability of reaching *q* = 0.05 was mapped onto this calibration curve by linear interpolation between adjacent points. The corresponding hard-selection coefficient was defined as the effective selection coefficient, *s_e_*.

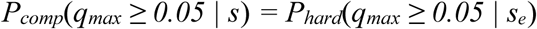

Next, we calculated |*s_e_*|/|*s*|. Values below one indicate attenuation of the assigned genotype-level effect relative to hard selection, whereas values above one indicate amplification.

Uncertainty in |*s_e_*| was estimated by non-parametric bootstrap resampling of simulation replicates. For each of 2000 bootstrap iterations, replicates were resampled with replacement independently within each hard-selection coefficient and competitive treatment. The hard-selection calibration was then refitted and |*s_e_*| re-estimated by linear interpolation. The 2.5^th^ and 97.5^th^ percentiles of the resulting bootstrap distribution were used as the 95% confidence interval for |*s_e_*|. Confidence intervals for |*s_e_*|/|*s*| were calculated from the corresponding bootstrap estimates.

To assess whether the calibration depended strongly on the choice of *q* = 0.05, we also compared the distributions of the maximum allele frequency (*q*_max_) reached by mutations under competitive selection with those under the matched hard-selection coefficient. We repeated this comparison across dominance coefficients *h* = 0, 0.1, 0.2 and 0.5 (Supplementary Fig. S3). We additionally provide SLiM and R code that allows users to generate a matched hard-selection reference under user-defined population parameters and estimate sₑ for alternative selection regimes from their mutation-frequency trajectories.

### 4.8 Reproductive variance and effective population size

Competitive selection can alter *N_e_*through both demographic effects on the number of breeders and increased variance in reproductive success. For each cohort we therefore recorded the number of gametic contributions made by every potential breeder, including zeros, and calculated the variance in reproductive success, *V_k_*. A short-term variance effective population size was estimated using the standard variance effective-size approximation (Wang et al. 2016):

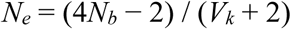

where *N_b_* is the number of potential breeders. These estimates were used to quantify differences in *Nₑ* among treatments and to guide the hard-selection sensitivity analysis described above.

### 4.9 Data analysis

Simulation output was processed in R. Threshold-crossing fractions were calculated relative to the number of newly arising offspring mutations registered in each replicate and then summarised across replicates. The full absolute-selection *q* = 0.05 calibration curve was used to estimate |*sₑ*| for each combination of assigned |*s*|, *D* and *R*. We examined both the mapping from |*s*| to |*sₑ*| and the ratio |*s_e_*|/|*s*|. Treatment effects were evaluated from independent replicate-level summaries rather than treating individual mutations as independent replicates.

## Data availability

All data analysed in this study were generated by the simulations described in the Methods. Source data underlying the main and Supplementary figures, including the simulation summaries used to estimate effective selection coefficients, genetic load and allele-frequency distributions, are provided with this paper. No empirical datasets were generated or analysed in this study.

## Code availability

The SLiM scripts and associated analysis code used for the population-genetic simulations are available from Thomas Birley’s GitHub repository: https://github.com/TBirley/Effective_s_calibration_SLiM. The repository also includes a reusable calibration workflow that generates hard-selection reference curves for user-defined population parameters and estimates the corresponding effective selection coefficient, sₑ, for alternative SLiM selection models. Code used for the repeated-contest and environmental-noise analyses is available from Jack Marsters’ GitHub repository: https://github.com/OrangeJack-et/multi_contest_sim.

## Author Contributions

C.v.O. developed the study concept and drafted the manuscript. J.N.M. and T.B. developed and performed the modelling and simulations and reviewed the manuscript. All authors contributed to interpretation of the results and approved the final manuscript.

## Use of generative AI

ChatGPT (OpenAI) was used during manuscript preparation to assist with checking and improving English-language expression, clarity and readability.

## References

Agrawal A.F., Whitlock M.C., 2012. Mutation load: the fitness of individuals in populations where deleterious alleles are abundant. Annual Review of Ecology, Evolution, and Systematics 43: 115–135. 10.1146/annurev-ecolsys-110411-160257

Almbro M., Simmons L.W., 2014. Sexual selection can remove an experimentally induced mutation load. Evolution 68: 295–300. 10.1111/evo.12238

Arbuthnott D., Rundle H.D., 2012. Sexual selection is ineffectual or inhibits the purging of deleterious mutations in Drosophila melanogaster. Evolution 66: 2127–2137. 10.1111/j.1558-5646.2012.01584.x

Arnold S.J., Wade M.J., 1984. On the measurement of natural and sexual selection: applications. Evolution 38: 720–734. 10.1111/j.1558-5646.1984.tb00345.x

Bell D.A., Kovach R.P., Robinson Z.L., Whiteley A.R., Reed T.E., 2021. The ecological causes and consequences of hard and soft selection. Ecology Letters 24: 1505–1521. 10.1111/ele.13754

Bertorelle, G., Raffini, F., Bosse, M., Bortoluzzi, C., Iannucci, A., Trucchi, E., Morales, H.E. and van Oosterhout, C., 2022. Genetic load: genomic estimates and applications in non-model animals. Nature Reviews Genetics, 23(8), pp.492–503. 10.1038/s41576-022-00448-x

Cally J.G., Stuart-Fox D., Holman L., 2019. Meta-analytic evidence that sexual selection improves population fitness. Nature Communications 10: 2017. 10.1038/s41467-019-10074-7

Cavill, E.L., Morales, H.E., Sun, X., Westbury, M.V., van Oosterhout, C., Accouche, W., Zora, A., Schulze, M.J., Shah, N., Adam, P. and Brooke, M.D.L., 2024. When birds of a feather flock together: Severe genomic erosion and the implications for genetic rescue in an endangered island passerine. Evolutionary Applications, 17 (7) 10.1111/eva.13739

Charlesworth B., 2009. Effective population size and patterns of molecular evolution and variation. Nature Reviews Genetics 10: 195–205. 10.1038/nrg2526

Charlesworth B., 2013. Why we are not dead one hundred times over. Evolution 67: 3354–3361. 10.1111/evo.12195

Chen, R.S., Soulsbury, C.D., Hench, K., van Oers, K. and Hoffman, J.I., 2025. Predicted deleterious mutations reveal the genetic architecture of male reproductive success in a lekking bird. Nature Ecology & Evolution, 9(10), pp.1924–1937. 10.1038/s41559-025-02802-8

Cherry J.L., Wakeley J., 2003. A diffusion approximation for selection and drift in a subdivided population. Genetics 163: 421–428. 10.1093/genetics/163.1.421

de Jong M.J., van Oosterhout C., Hoelzel A.R., Janke A., 2024. Moderating the neutralist-selectionist debate: exactly which propositions are we debating, and which arguments are valid? Biological Reviews 99: 23–55. 10.1111/brv.13010

Dussex N., Morales H.E., Grossen C., Dalén L., van Oosterhout C., 2023. Purging and accumulation of genetic load in conservation. Trends in Ecology & Evolution 38: 961–969. 10.1016/j.tree.2023.05.008

Fontsere, C., Speak, S.A., Caven, A.J., Rodríguez, J.A., Wang, X., Pacheco, C., Cassatt-Johnstone, M., Femerling, G., Maloney, B., Balacco, J. and Collins, J., 2025. Persistent genomic erosion in whooping cranes despite demographic recovery. Molecular Ecology, 34(23), p.e70088. 10.1111/mec.70088

Haller B.C., Messer P.W., 2023. SLiM 4: Multispecies eco-evolutionary modeling. The American Naturalist 201: E127–E139. 10.1086/723601

Hsu Y., Earley R.L., Wolf L.L., 2006. Modulation of aggressive behaviour by fighting experience: mechanisms and contest outcomes. Biological Reviews 81: 33–74. 10.1017/S146479310500686X

Hunt J., Breuker C.J., Sadowski J.A., Moore A.J., 2009. Male–male competition, female mate choice and their interaction: determining total sexual selection. Journal of Evolutionary Biology 22: 13–26. 10.1111/j.1420-9101.2008.01633.x

Lesecque Y., Keightley P.D., Eyre-Walker A., 2012. A resolution of the mutation load paradox in humans. Genetics 191: 1321–1330. 10.1534/genetics.112.140343

López-Cortegano E., Charlesworth B., 2026. The effect of a reduction in population size on mean fitness and inbreeding depression. bioRxiv. 10.64898/2026.05.15.725556

Lumley, A.J., Michalczyk, Ł., Kitson, J.J., Spurgin, L.G., Morrison, C.A., Godwin, J.L., Dickinson, M.E., Martin, O.Y., Emerson, B.C., Chapman, T. and Gage, M.J., 2015. Sexual selection protects against extinction. Nature, 522(7557), pp.470–473. 10.1038/nature14419

Marrec L., 2026. Impact of genotype-dependent dispersal on mutation fixation in subdivided populations. Journal of Evolutionary Biology 39: 262–271. 10.1093/jeb/voaf134

McGlothlin J.W., 2010. Combining selective episodes to estimate lifetime nonlinear selection. Evolution 64: 1377–1385. 10.1111/j.1558-5646.2009.00914.x

McGuigan K., Petfield D., Blows M.W., 2011. Reducing mutation load through sexual selection on males. Evolution 65: 2816–2829. 10.1111/j.1558-5646.2011.01346.x

Noel, E., Fruitet, E., Lelaurin, D., Bonel, N., Segard, A., Sarda, V., Jarne, P. and David, P., 2019. Sexual selection and inbreeding: Two efficient ways to limit the accumulation of deleterious mutations. Evolution Letters, 3(1), pp.80–92. 10.1002/evl3.93

Ohta T., 1972. Fixation probability of a mutant influenced by random fluctuation of selection intensity. Genetical Research 19: 33–38. 10.1017/S0016672300014245

Ohta T., 1973. Slightly deleterious mutant substitutions in evolution. Nature 246: 96–98. 10.1038/246096a0

Pál C., Papp B., Hurst L.D., 2001. Highly expressed genes in yeast evolve slowly. Genetics 158: 927–931. 10.1093/genetics/158.2.927

Pointer, M.D., Nash, W.J., Gage, M.J., Chapman, T., Maklakov, A.A. and Richardson, D.S., 2026. Sexual selection purges mutation load, but not overall genetic diversity, decreasing vulnerability to extinction. Proceedings of the National Academy of Sciences, 123(37), p.e2608675123. 10.1073/pnas.2608675123

Reznick D., 2016. Hard and soft selection revisited: how evolution by natural selection works in the real world. Journal of Heredity 107: 3–14. 10.1093/jhered/esv076

Rowe L., Houle D., 1996. The lek paradox and the capture of genetic variance by condition dependent traits. Proceedings of the Royal Society B 263: 1415–1421. 10.1098/rspb.1996.0207

Siller S., 2001. Sexual selection and the maintenance of sex. Nature 411: 689–692. 10.1038/35079578

Speak, S.A., Birley, T., Bortoluzzi, C., Clark, M.D., Percival-Alwyn, L., Morales, H.E. and van Oosterhout, C., 2024. Genomics-informed captive breeding can reduce inbreeding depression and the genetic load in zoo populations. Molecular Ecology Resources, 24(7), p.e13967. 10.1111/1755-0998.13967

Thurman T.J., Barrett R.D.H., 2016. The genetic consequences of selection in natural populations. Molecular Ecology 25: 1429–1448. 10.1111/mec.13559

Trucchi, E., Massa, P., Giannelli, F., Latrille, T., Gargano, M., Nitta Fernandes, F.A., Ancona, L., Stenseth, N.C., Ferrer Obiol, J., Paris, J. and Bertorelle, G., 2025. High gene expression predicts extremely low segregation of deleterious mutations in large penguin populations. Molecular biology and evolution, 42(6), p.msaf146. 10.1093/molbev/msaf146

Tschol M., Reid J.M., Bocedi G., 2023. Environmental variance in male mating success modulates the positive versus negative impacts of sexual selection on genetic load. Journal of Evolutionary Biology 36: 1242–1254. 10.1111/jeb.14202

van Oosterhout, C., Helsen, P., Fienieg, E., Fontsere, C., Rasmussen, L., Bosse, M., Pereboom, Z., Hvilsom, C. and Morales, H.E., 2026. Genomics-informed management of ex situ populations. Conservation Biology, p.e70360. 10.1111/cobi.70360

van Oosterhout C., Marcu D., Immler S., 2022. Accounting for the genetic load in assisted reproductive technology. Clinical and Translational Medicine 12: e864. 10.1002/ctm2.864

Wallace B., 1975. Hard and soft selection revisited. Evolution 29: 465–473. 10.1111/j.1558-5646.1975.tb00836.x

Wanders, K., Chen, G., Feng, S., Zhang, G., Székely, T., Bruford, M., Végvári, Z., Eichhorn, G. and Urrutia, A., 2023. Polygamy and purifying selection in birds. Evolution, 77(1), pp.276–288. 10.1093/evolut/qpac010

Wang J., Santiago E., Caballero A., 2016. Prediction and estimation of effective population size. Heredity 117: 193–206. 10.1038/hdy.2016.43

Wang, X., Stuart, A., Norris, K., Henshaw, S., Strang, A., Pacheco, C., Nielsen, S.D., Waite, M., Hume, J.P., Ruhomaun, K. and Brace, S., 2026. Conservation rescued the Mauritius kestrel from extinction but not from genomic erosion. Biorxiv, pp.2026–06. 10.64898/2026.06.23.733933

Wang, X., Fontsere, C., Alva Caballero, L.X., Nielsen, S.D., Groombridge, J.J., Hansson, B., van Oosterhout, C., Pacheco, C. and Morales, H.E., 2026b. Genomic erosion across avian lineages in the context of their evolutionary history. Molecular biology and evolution, 43(4), p.msag070.. 10.1093/molbev/msag070

Whitlock M.C., 2000. Fixation of new alleles and the extinction of small populations: drift load, beneficial alleles, and sexual selection. Evolution 54(6): 1855–1861. 10.1111/j.0014-3820.2000.tb01232.x

Whitlock M.C., Agrawal A.F., 2009. Purging the genome with sexual selection: reducing mutation load through selection on males. Evolution 63: 569–582. 10.1111/j.1558-5646.2008.00558.x

Wu, Z., Cai, X., Zhang, X., Liu, Y., Tian, G.B., Yang, J.R. and Chen, X., 2022. Expression level is a major modifier of the fitness landscape of a protein coding gene. Nature Ecology & Evolution, 6(1), pp.103–115.. 10.1038/s41559-021-01578-x

