## Supplementary information for "Reproductive competition amplifies purifying selection and reduces genetic load"

The supplementary analyses first test the robustness of the effective-selection calibration to differences in drift, allele-frequency threshold, dominance, candidate population size, mutation rate and recombination parameter ρ. They then examine the contribution of reproductive skew, how repeated contests improve competitive discrimination under transient environmental variation, and the empirical basis of the reproductive-skew parameterisation.

### S1. Effective selection calibration and robustness

#### S1.1 Effective population size and neutral threshold crossing

Calibration of $s_e$ depends on the probability that a mutation reaches a specified frequency under the absolute-selection reference. Because differences in $N_e$ alter the strength of drift, we tested whether residual differences in $N_e$ could affect the calibration. We varied carrying capacity ($K$) in the absolute-selection model and compared both realised $N_e$ and the probability that neutral mutations reached $q = 0.05$ with the competitive treatment.

Changing $K$ altered $N_e$, but had little effect on the probability that neutral mutations reached $q = 0.05$ (Fig. S1). The competitive treatment showed a similar neutral threshold-crossing probability. Thus, the much larger treatment differences in $s_e$ are unlikely to result simply from differences in drift.

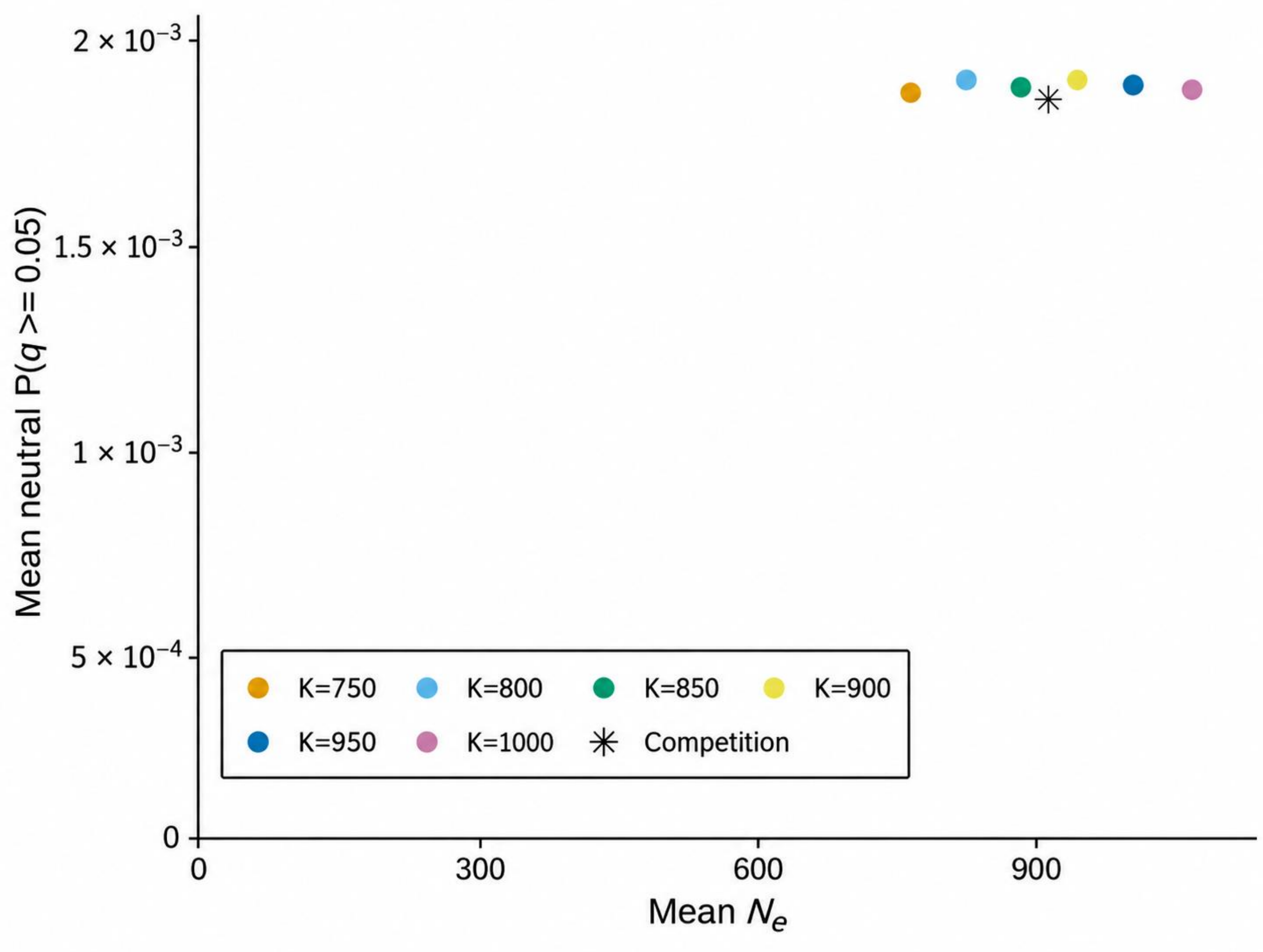


**Figure S1.** Matching drift between absolute and competitive selection. Mean effective population size and the probability that neutral mutations reached $q = 0.05$ under competitive selection and alternative absolute-selection values of $K$. Changing $K$ altered $N_e$ but had little effect on neutral threshold crossing, and the competitive treatment fell within a narrow range.

**S1.2 Robustness of the effective-selection calibration across allele frequencies**

The effective selection coefficient was calibrated using the probability that a newly arisen mutation reached $q = 0.05$. To test whether this single threshold provided a representative mapping between competitive and hard selection, we compared the full distributions of the maximum allele frequency ($q_{max}$) reached by mutations under the two selection regimes. For each competitive regime, the corresponding hard-selection coefficient was first identified from the $q = 0.05$ calibration, after which the full $q_{max}$ distributions were compared.

For codominant mutations ($h = 0.5$) with assigned $s = -0.0025$, the competitive treatments and their corresponding hard-selection references showed close agreement across the $q_{max}$

distribution (Fig. S2). These competitive regimes encompassed both attenuation and strong amplification of the effective selection coefficient. Thus, for the codominant mutations analysed here, calibration at $q = 0.05$ captured a broader equivalence in mutation fate. For the other assigned $s$ values ($s = 0$, −0.001 and −0.005), competitive treatments likewise closely matched their corresponding hard-selection references across the $q_{max}$ distributions. We additionally tested whether this correspondence was sensitive to dominance by comparing hard and competitive selection across $h = 0$, 0.1, 0.2 and 0.5. Although increasing $h$ reduced the probability of reaching $q$=0.05, the $q_{max}$ distributions and threshold-crossing probabilities remained closely matched between selection modes across dominance coefficients (Fig. S3). This shows that the correspondence was not specific to the codominant mutations used in the main simulations.

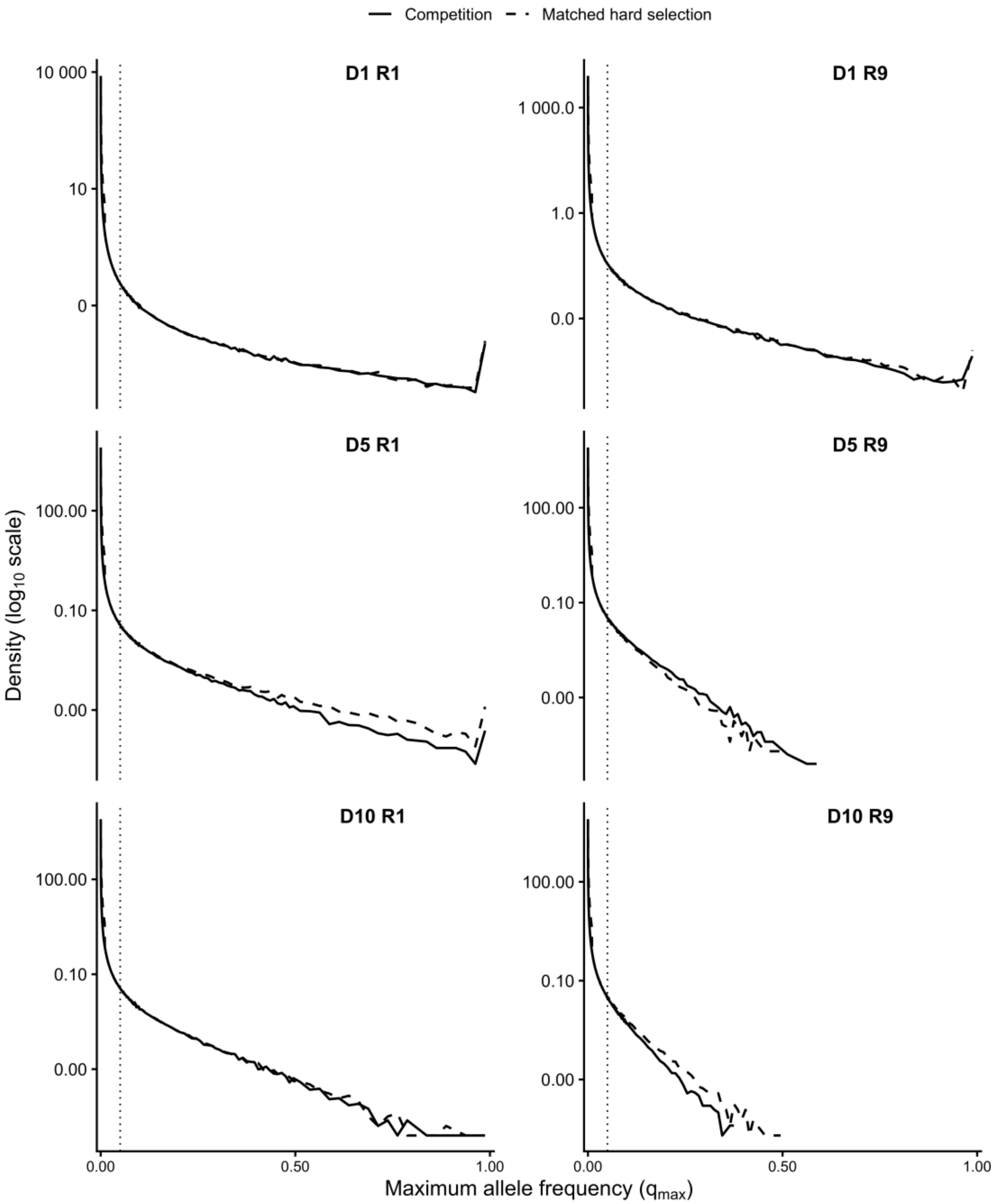


**Figure S2.** Robustness of the effective-selection calibration across allele frequencies. Distributions of the maximum allele frequency ($q_{max}$) reached by deleterious mutations under each competitive regime (solid lines) and its corresponding hard-selection reference (dashed lines), calibrated at q = 0.05. The vertical dashed line marks $q = 0.05$. *D* denotes distinct rivals and *R* number of contest rounds per rival. Density is shown on a logarithmic scale; y-axis ranges vary among panels.

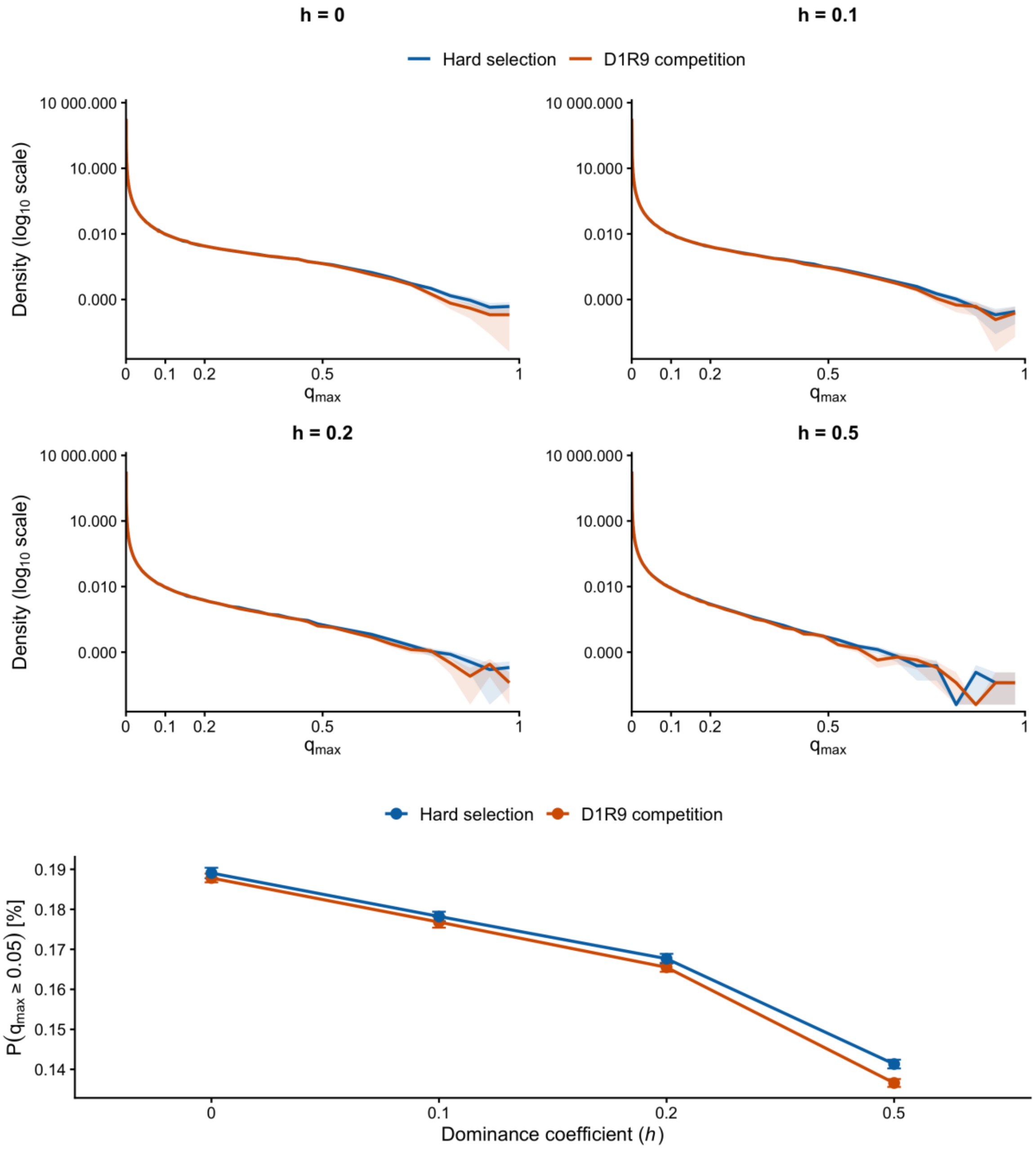


Figure S3. Robustness of mutation trajectories to dominance. Maximum allele-frequency ($q_{max}$) distributions under hard selection and *D*1 *R*9 competition for deleterious mutations with dominance coefficients $h$ = 0, 0.1, 0.2, and 0.5. Lines show mean distributions across replicate simulations and shaded regions show 95% confidence intervals; density is plotted on a log10 scale. The lower plot shows the probability that newly arisen mutations reached $q_{max} \geq 0.05$ under hard and competitive selection across dominance coefficients. Increasing $h$ reduced threshold-crossing probability, but the close correspondence between selection modes was maintained across dominance coefficients.

#### S1.3 Relative efficiency of competitive selection

We also expressed effective selection relative to the assigned genotype-level effect as $|s_e|/|s|$. This ratio was broadly stable across the tested values of $|s|$ within each competitive regime (Fig. S4), supporting the interpretation that selection mode acts approximately as a multiplier of genotype-level fitness effects over this range. Values below one indicate attenuation relative to absolute selection, whereas values above one indicate amplification. The next analysis isolates the contest-level mechanism that produces this increase in discrimination.

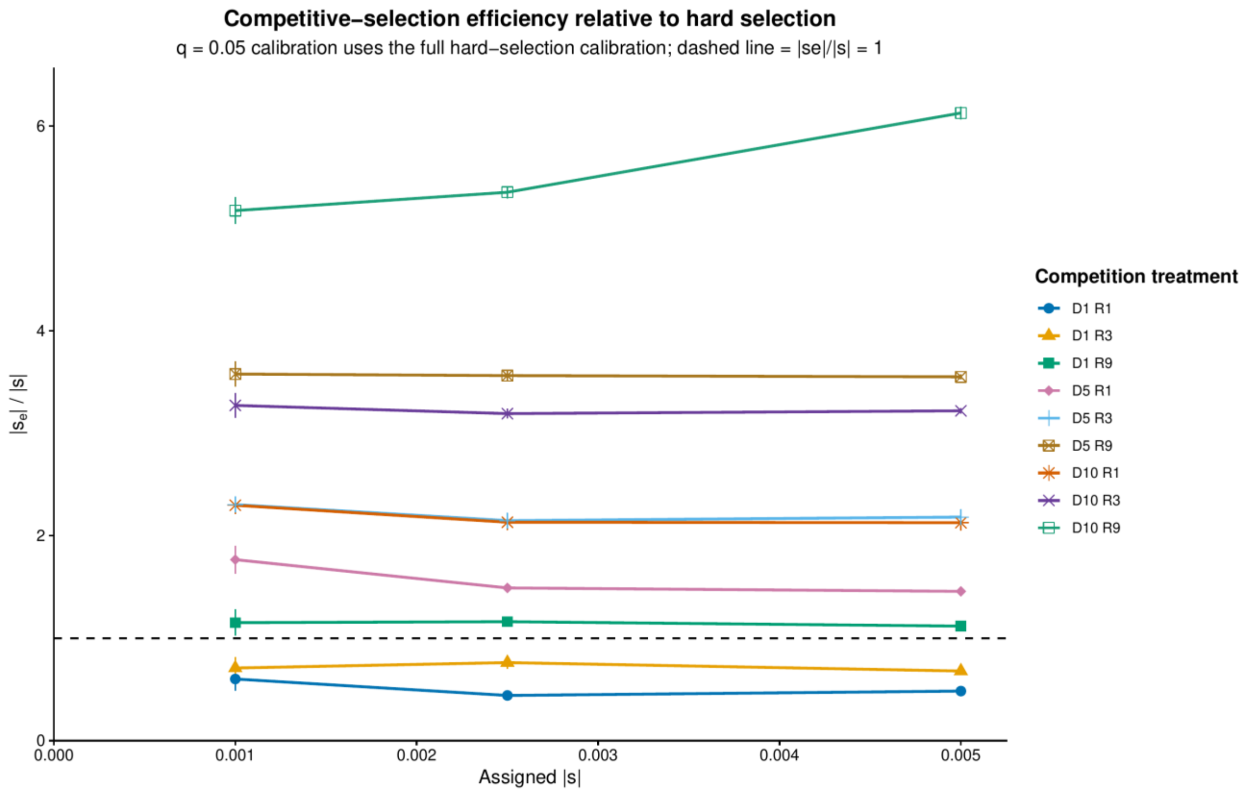


Figure S4. Competitive-selection efficiency relative to absolute selection. The ratio $|s_e|/|s|$ was calculated from the $q = 0.05$ calibration shown in main-text Figure 1. The dashed line denotes equal efficiency ($|s_e|/|s| = 1$). Repeated contests against a single rival progressively increased effective selection, whereas exposure to multiple distinct rivals produced substantially stronger amplification, especially when combined with repeated rounds.

#### S1.4 Sensitivity of selection amplification to dominance

To test whether selection amplification depended on dominance, we analysed six dominance coefficients ($h$ = 0, 0.01, 0.05, 0.1, 0.2 and 0.5) and two assigned deleterious effects ($s = -0.0025$

and −0.005). Hard selection was compared with three competitive regimes (R1/D1, R3/D5 and R9/D10), using 100 replicates per treatment. For each *h*, the effective coefficient $s_e$ was calibrated against the probability of a new mutation reaching $q \geq 0.05$ under hard selection. Amplification was expressed as $|s_e|/|s|$. Confidence intervals were obtained from 2,000 replicate-level bootstrap resamples, recalibrating the hard-selection reference in each iteration.

The qualitative ordering of competition regimes was broadly conserved across dominance coefficients (Fig. S5). Weak competition generally attenuated purifying selection relative to hard selection, whereas intermediate and strong competition amplified it. Under intermediate competition, relative amplification tended to decline with increasing *h*, even though threshold-crossing probabilities decreased with *h* in all regimes. Estimates were less precise for strongly recessive mutations, whose heterozygous effects were masked; some $h = 0$ calibrations were not identifiable within the hard-selection reference range. Thus, the qualitative conclusion does not depend on assuming $h = 0.5$, although the magnitude and identifiability of $|s_e|/|s|$ depend on dominance.

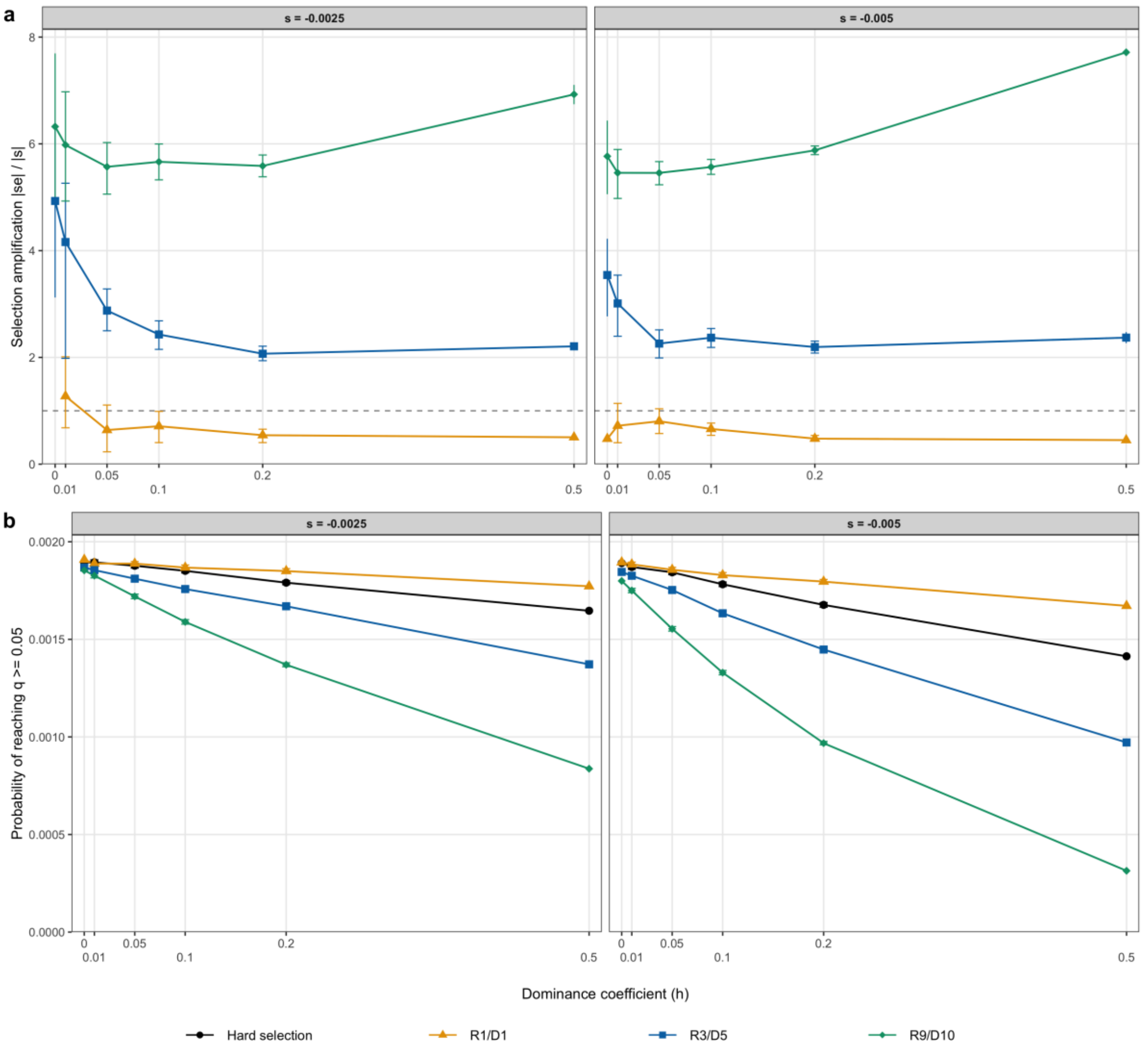


**Figure S5.** Influence of dominance on competitive-selection effectiveness. (a) Amplification $|s_e|/|s|$ across $h = 0$, 0.01, 0.05, 0.1, 0.2 and 0.5, for assigned $s = -0.0025$ and $-0.005$. Values above one indicate stronger effective purifying selection than hard selection; error bars indicate 95% bootstrap confidence intervals where at least 95% of iterations yielded an identifiable calibration. (b) Probability that a newly arising deleterious mutation reached $q \geq 0.05$ under hard selection and R1/D1, R3/D5 and R9/D10 competition. Each treatment comprised 100 replicates.

### S1.5 Effective heterozygous selection across dominance coefficients

To express selection in terms of the effect experienced by rare heterozygotes, we compared the assigned heterozygous effect $|hs|$ with the calibrated effective effect $|hs_e|$, using the same dominance coefficient in the hard-selection reference. The analysis included $s = -0.0025$ and $-0.005$ and $h = 0.05$, 0.1, 0.2 and 0.5. The $h = 0$ case has no heterozygous effect, whereas calibration at h = 0.01 was insufficiently informative. Uncertainty was propagated from 2,000 bootstrap iterations.

Effective heterozygous selection increased with assigned |*hs*|, but depended strongly on competition intensity (Fig. S6). A proportional fit, $|hs_e| = \beta|hs|$, yielded $\beta = 0.47$ (95% bootstrap CI 0.45–0.50) under R1/D1, 2.32 (2.24–2.39) under R3/D5 and 7.24 (7.19–7.30) under R9/D10. The stronger regimes showed positive curvature, so these fitted slopes summarize the analysed range rather than imply a constant multiplier. Because $|hs_e|/|hs| = |s_e|/|s|$ for $h > 0$, this analysis provides a heterozygote-scale representation of the amplification estimates rather than an independent calibration.

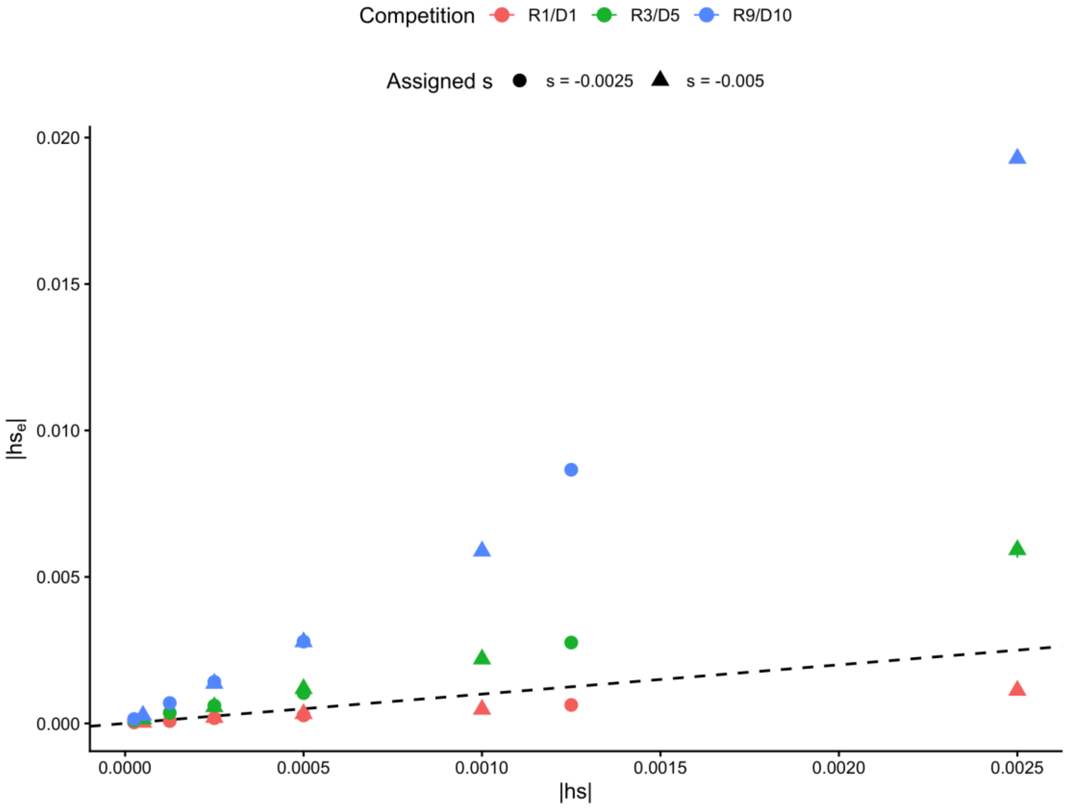


**Figure S6.** Assigned versus effective heterozygous selection under different competitive regimes. Assigned |*hs*| is plotted against calibrated $|hs_e|$ for s = −0.0025 (circles) and −0.005 (triangles), with $h$ = 0.05, 0.1, 0.2 and 0.5. Colours distinguish R1/D1, R3/D5 and R9/D10; the dashed line is the 1:1 reference.

#### S1.6 Candidate population size and neutral calibration

We tested the sensitivity of effective-selection estimates to candidate population size ($N_c$ = 100, 250, 500, 1,000, 2,500, 5,000 and 25,000), using 100 replicates per treatment and matching ($N_c$)

between hard and competitive selection. At $N_c = 500$, estimates of $|s_e|$ were substantially larger than those obtained at greater population sizes (Fig. S7a,c). The neutral threshold-crossing probability was approximately 4.2% lower under competition than hard selection at $N_c = 500$, compared with 0.6% at $N_c = 5000$ (Table S1). Normalising threshold-crossing probabilities by their respective neutral baselines substantially reduced the apparent amplification at $N_c = 500$, whereas the adjustment had little effect at $N_c = 5000$ (Fig. S7a,b).

Following neutral normalisation, estimates of $|s_e|$ were broadly consistent across $N_c \geq 500$, with the greatest variation occurring at $N_c = 500$ and estimates converging towards similar values at larger population sizes (Fig. S7c). At $N_c = 100$ and 250, calibration was less reliable, with greater uncertainty and inconsistent estimates of selection amplification. At these small effective population sizes ($N_e \approx 20$–55), $2N_e|s| < 1$ for the assigned deleterious effects used in the competitive treatments, so genetic drift dominates allele-frequency changes. Consequently, threshold-crossing probabilities become relatively insensitive to differences in selection strength, making se difficult to estimate precisely. This limitation of the calibration does not imply that competitive selection ceases to amplify purifying selection in small populations. Rather, it shows that genetic drift obscures differences in selection strength, reducing the precision of our calibration. Estimates of $N_e/N_c$ remained broadly consistent across population sizes, averaging approximately 0.22 under hard selection and 0.20 under competition, with some deviations at the smallest population sizes (Table S2). Overall, these results indicate that effective-selection estimates are reasonably robust to candidate population size at $N_c \geq 500$, but become less reliable in very small, drift-dominated populations.

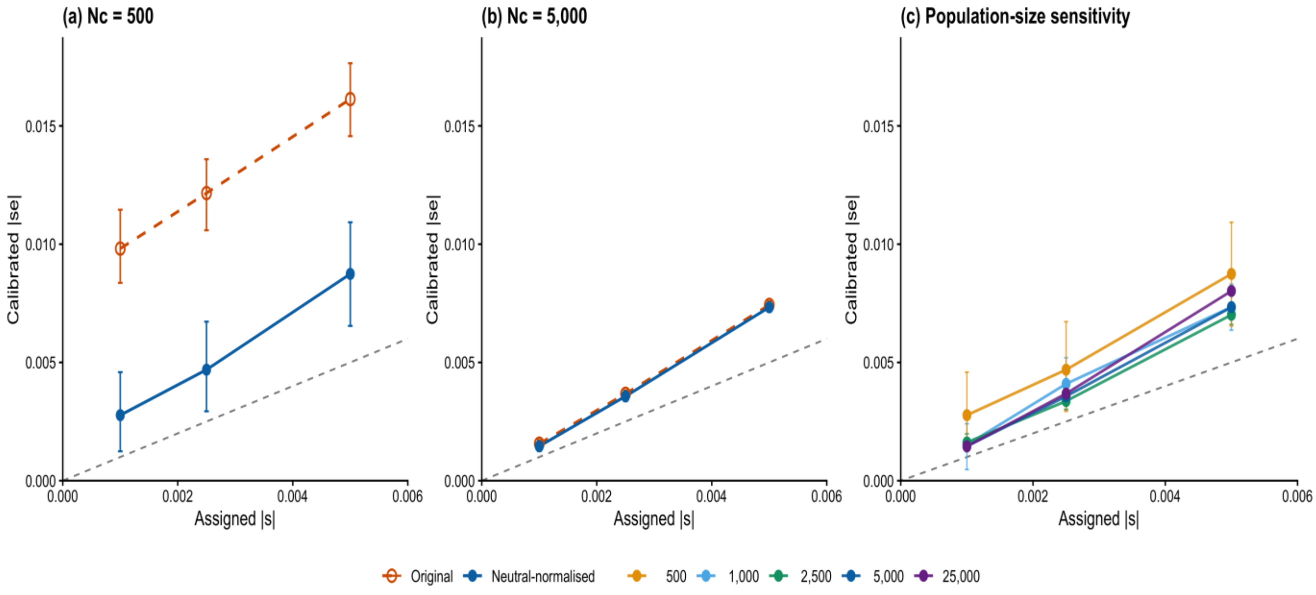


**Figure S7. Robustness of effective-selection estimates across candidate population sizes and neutral calibration.** Relationship between assigned selection coefficients (|*s*|) and calibrated effective selection coefficients ($|s_e|$). (a) Original and neutral-normalised estimates for $N_c$ = 500. (b) Equivalent comparison for $N_c$ = 5,000. (c) Comparison of effective-selection estimates across $N_c$ = 500, 1,000, 2,500, 5,000 and 25,000. Error bars indicate 95% bootstrap confidence intervals; the grey dashed diagonal represents ($|s_e|$ = |*s*|).

**Table S1.** Neutral threshold-crossing probabilities under hard and competitive selection.

| $N_c$ | **Hard selection $P_0$** | **Competition $P_0$** | $P_{0,Comp}/ P_{0,HS}$ |
|---|---|---|---|
| 100 | 0.086744 | 0.072548 | 0.836 |
| 250 | 0.035354 | 0.031952 | 0.904 |
| 500 | 0.017865 | 0.017116 | 0.958 |
| 1000 | 0.009067 | 0.008840 | 0.975 |
| 2500 | 0.003776 | 0.003709 | 0.982 |
| 5000 | 0.001907 | 0.001896 | 0.994 |

The neutral probability $P_0$ is the pooled fraction of newly tracked neutral mutations reaching $q \geq$ 0.05 across 100 replicates per treatment. The ratio $P_{0,Comp}/ P_{0,HS}$ indicates the difference in neutral baselines between regimes.

**Table S2.** Effective population size estimated from variance in reproductive success.

| Selection mode | $N_c$ | Mean $N_e$ | $N_e/N_c$ |
|---|---|---|---|
| Hard | 100 | 21.53 | 0.2153 |
| Competition | 100 | 22.19 | 0.2219 |
| Hard | 250 | 56.18 | 0.2247 |
| Competition | 250 | 52.18 | 0.2087 |
| Hard | 500 | 111.66 | 0.2233 |
| Competition | 500 | 101.30 | 0.2026 |
| Hard | 1,000 | 222.87 | 0.2229 |
| Competition | 1,000 | 199.68 | 0.1997 |
| Hard | 2,500 | 555.97 | 0.2224 |
| Competition | 2,500 | 495.20 | 0.1981 |
| Hard | 5,000 | 1,111.38 | 0.2223 |
| Competition | 5,000 | 987.17 | 0.1974 |
| Hard | 25,000 | 5,555.54 | 0.2222 |
| Competition | 25,000 | 4,921.36 | 0.1969 |

$N_c$ is the number of candidate offspring before selection. Mean $N_e$ was calculated using the variance in reproductive success ($V_k$) method across 100 replicates per parameter combination, with values subsequently averaged across assigned selection coefficients within each selection regime.

#### S1.7 Sensitivity to mutation rate

To assess whether estimates of effective selection depended on mutation rate, we repeated R1/D5 competition across $\mu = 10^{-6}$, $10^{-5}$ and $10^{-4}$, using a separate hard-selection calibration for each. Amplification $|s_e|/|s|$ ranged from 1.469 to 1.668 (Table S3). The estimates at $\mu = 10^{-5}$ and $10^{-4}$ were nearly identical, whereas $\mu = 10^{-6}$ gave a somewhat higher and less precise estimate. Thus, selection amplification persisted across a 100-fold range in mutation rate.

**Table S3.** Sensitivity of effective-selection estimates to mutation rate.

| **Mutation rate ($\mu$)** | **P($q \geq 0.05$)** | **$\|s_e\|$** | **$\|s_e\|/\|s\|$ (95% CI)** |
|---|---|---|---|
| 10-6 | 0.001511 | 0.004169 | 1.668 (1.456-1.905) |
| 10-5 | 0.001537 | 0.003672 | 1.469 (1.409-1.525) |
| 10-4 | 0.001539 | 0.003677 | 1.471 (1.452-1.489) |

R1/D5 competition used assigned $s = -0.0025$. For each $\mu$, $|s_e|$ was obtained by matching the probability of a new deleterious mutation reaching $q \geq 0.05$ to its independently generated hard-selection calibration. Amplification is $|s_e|/|s|$. Results are based on 100 replicates per treatment and 2,000 bootstrap iterations.

### S1.8 Sensitivity to recombination parameter ρ

To assess sensitivity to recombination parameter ρ, we repeated R1/D5 competition at $\rho = 0.005$, 0.05 and 0.5, using a separate hard-selection calibration for each. Estimated amplification $|s_e|/|s|$ was broadly consistent, ranging from 1.436 to 1.469, with substantially overlapping 95% bootstrap confidence intervals (Table S4). This supports robustness of selection amplification across the tested 100-fold variation in ρ, without establishing exact equivalence among treatments.

**Table S4.** Sensitivity of effective-selection estimates to recombination parameter ρ.

| **Recombination rate ($\rho$)** | **P($q \geq 0.05$)** | **$\|s_e\|$** | **$\|s_e\|/\|s\|$ (95% CI)** |
|---|---|---|---|
| 0.005 | 0.001538 | 0.003595 | 1.438 (1.387–1.490) |
| 0.05 | 0.001546 | 0.003591 | 1.436 (1.380–1.488) |
| 0.5 | 0.001537 | 0.003672 | 1.469 (1.409-1.525) |

R1/D5 competition used assigned $s = -0.0025$, $K = 1{,}000$, $N_c = 5{,}000$, $\mu = 10^{-5}$ and $h = 0.5$. For each $\rho$, $|s_e|$ was obtained by matching the probability of a new deleterious mutation reaching $q \geq 0.05$ to an independent hard-selection calibration. Amplification is $|s_e|/|s|$. Results are based on 100 replicates per treatment and 2,000 bootstrap iterations.

#### S1.9 Random reproductive success control

To distinguish the effects of reproductive skew from those of competitive selection, we performed an additional control in which reproductive success was drawn from the same Gamma distribution used in the competition model but assigned randomly among individuals rather than according to competitive rank. The control used $s = -0.0025$, $h = 0.5$, $D = 10$, and $R = 9$, with all other simulation and mutation-tracking procedures unchanged. We compared this treatment with the corresponding competition-linked treatment and a neutral treatment ($s = 0$).

Random assignment of reproductive success produced mutation-frequency trajectories that were very similar to the neutral treatment across all frequency thresholds (Fig. S8). In contrast, when reproductive success was allocated according to competitive performance, substantially fewer mutations reached $q \geq 0.05$, with the difference becoming increasingly pronounced at higher frequencies. Very few mutations reached $q \geq 0.20$ under competition-linked reproductive success, and progression to $q \geq 0.50$ or fixation was effectively eliminated. Thus, reproductive skew alone did not generate the increased efficiency of selection observed under competition. Instead, the effect depended on the association between genotype-dependent competitive performance and reproductive success.

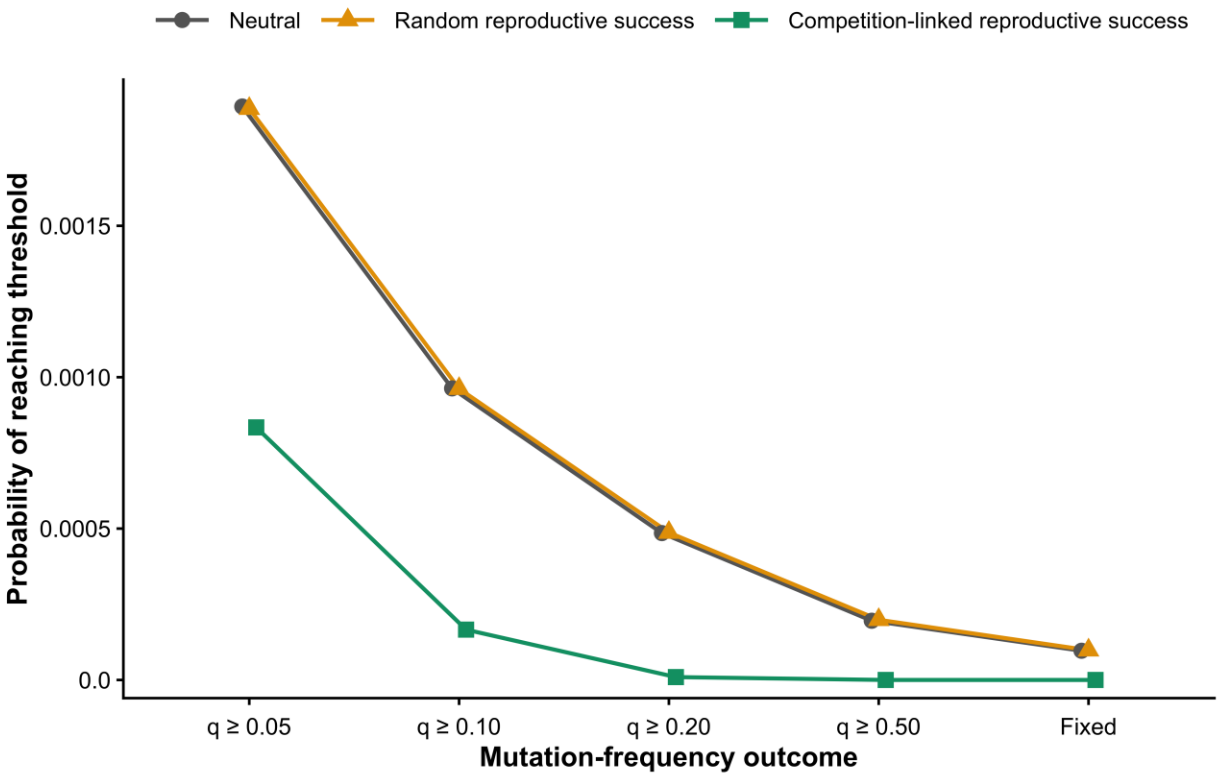


**Figure S8**. Random reproductive success control for the effect of reproductive skew on deleterious-mutation trajectories. Probability that newly arising mutations reached allele-frequency thresholds of $q \geq 0.05$, $q \geq 0.10$, $q \geq 0.20$, and $q \geq 0.50$, or fixation, under neutral, random reproductive success, and competition-linked reproductive success. For the latter two treatments, reproductive success followed the same Gamma distribution, but reproductive weights were either assigned randomly among individuals or allocated according to competitive rank. Both treatments used $s = -0.0025$, $h = 0.5$, $D = 10$, and $R = 9$; the neutral treatment used $s = 0$. Points show mean probabilities across simulation replicates. Random reproductive success closely reproduced the neutral mutation trajectory, whereas linking reproductive success to competitive performance strongly reduced the probability that deleterious mutations progressed to higher frequencies.

### S2. Why repeated contests improve competitive discrimination

To isolate the contest-level mechanism, we considered pairs of individuals with realised loads $L_A$ and $L_B$, with fitness $w = e^{-L}$. For a single contest, the probability that individual A wins is determined only by the difference in realised load, $\Delta L = L_B - L_A$:

$$P(\text{A wins}) = w_A/(w_A + w_B) = 1/(1 + e^{-\Delta L}).$$

For an odd number of independent contests, the overall winner is the individual winning the majority. Simulated probabilities closely followed the corresponding binomial majority-vote prediction without fitted parameters (Fig. S9), validating the repeated-contest implementation. As the number of contests increased, a persistent fitness difference was resolved with progressively greater reliability.

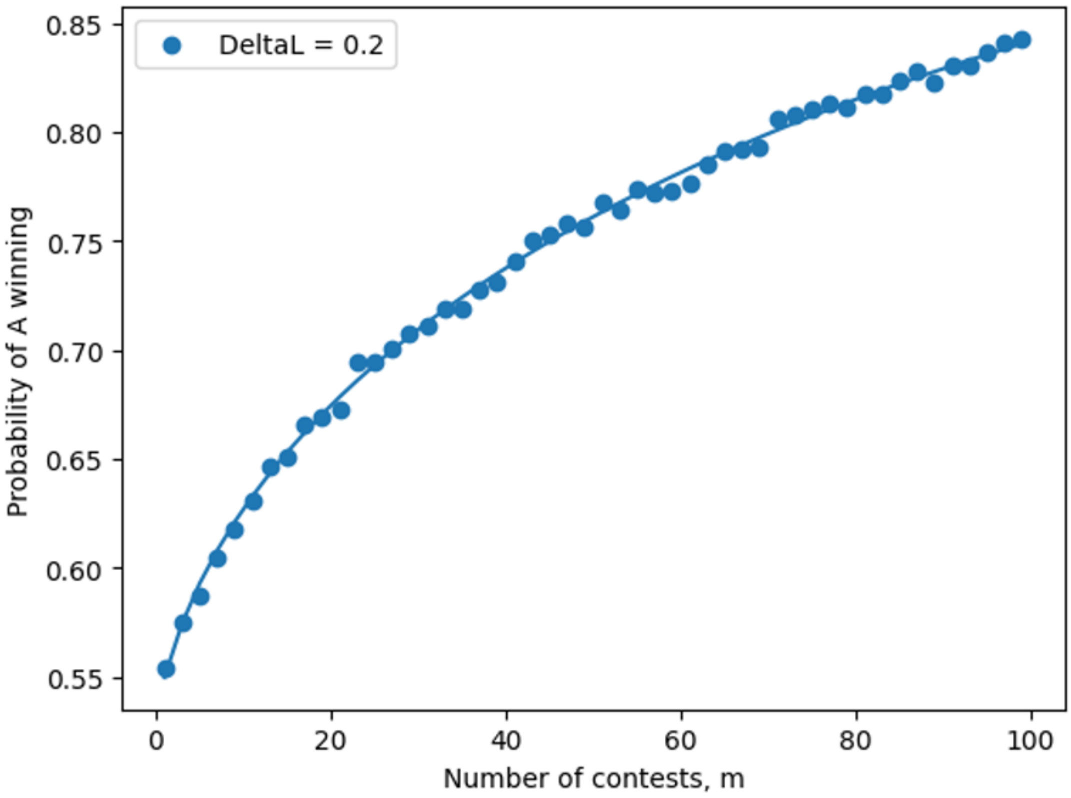


**Figure S9**. Validation of the repeated-contest simulation against the analytical majority-vote prediction. For a fixed difference in realised load of $\Delta L = 0.2$ and no environmental noise, points show simulated probabilities that the fitter individual wins the contest series as the number of contests increases. The solid curve is the binomial majority-vote prediction using the single-contest probability; no parameters were fitted.

#### S2.1 Transient environmental variation

To test whether repeated contests can recover persistent fitness differences when individual contests are noisy, we added a transient environmental deviation to realised load before each contest:

$$L_i' = L_i + \varepsilon_i, \quad \varepsilon_i \sim N(0, \sigma_E^2).$$

The deviation was drawn independently for each individual and each contest, representing temporary variation in competitive performance rather than a persistent environmental effect. Increasing $\sigma_E$ progressively reduced the probability that the fitter individual won a three-contest series, especially when the underlying difference in realised load ($\Delta L$) was small (Fig. S10). Larger underlying fitness differences remained detectable under greater environmental variation. The random-pair treatment remained at approximately 0.5, as expected when neither member of the pair was defined a priori as fitter.

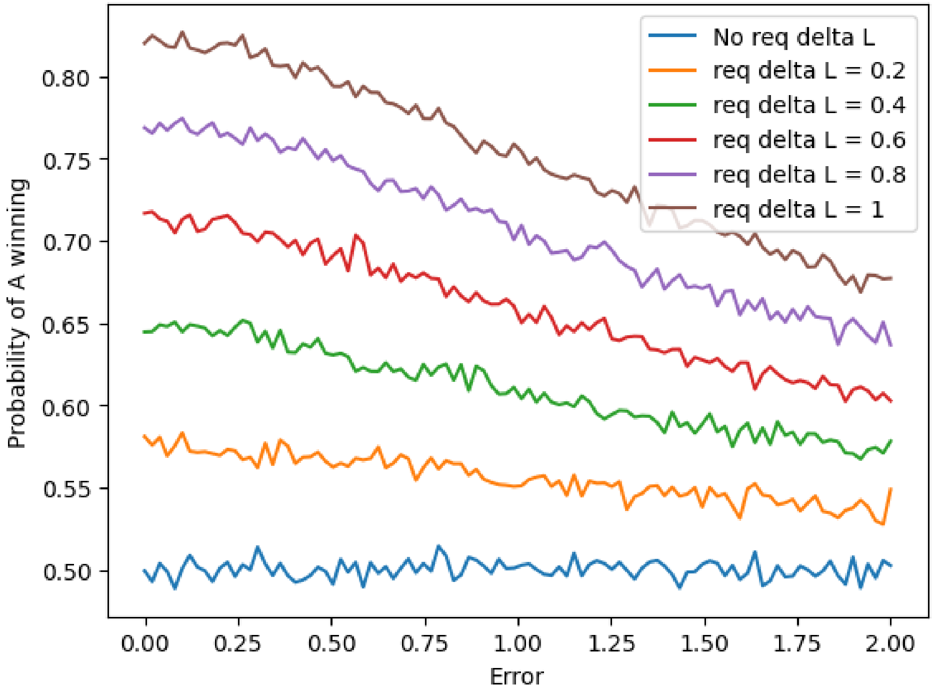


**Figure S10**. Environmental variation reduces the resolution of competitive assessment. Probability that the fitter individual wins a three-contest series as environmental variation ($\sigma_E$) increases, shown for different differences in realised load ($\Delta L$). Each environmental deviation was independently redrawn before every contest. Increasing environmental variation progressively obscured the underlying fitness difference, whereas larger $\Delta L$ remained detectable at higher levels of noise. The random-pair treatment remains at $P = 0.5$, as expected when neither individual is defined a priori as fitter.

Together, these contest-level analyses show that repeated assessment can convert a persistent fitness difference into increasingly reliable competitive success, while averaging over transient environmental variation. In the population simulations, this competitive ranking is translated into reproductive success using Gamma-distributed weights; their empirical calibration is described below.

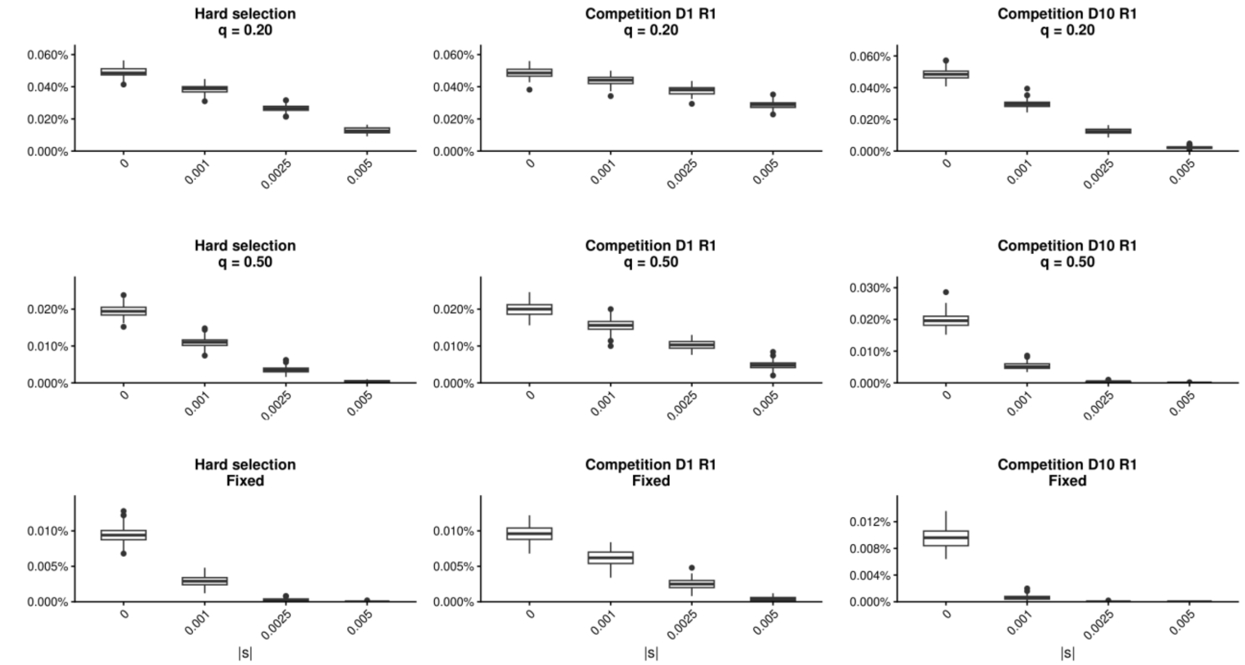


Figure S11. Competitive selection changes the probability that deleterious mutations reach high frequency. Fractions of newly arisen mutations reaching $q$=0.2, $q$=0.5, or fixation under absolute selection (left), competition with one rival and one round (D1 R1; middle), and competition with ten distinct rivals and one round per rival (D10 R1; right). Boxplots summarise replicate-level fractions across assigned $|s|$. Weak competition increased high-frequency persistence relative to absolute selection, whereas competition against ten distinct rivals strongly reduced high-frequency crossing and fixation.

### S3. Empirical calibration of Gamma-distributed reproductive output

Rationale. To model reproductive skew, we used Gamma-distributed relative reproductive weights. Most empirical studies do not fit Gamma distributions directly, but report the mean and variance in reproductive success, or the standardised variance $I = \mathrm{Var}(R)/\mathrm{E}(R)^2$ (the opportunity for selection). These values can be converted to a moment-matched Gamma distribution. For a Gamma distribution with shape α and scale θ, $\mathrm{E}(R) = \alpha\theta$ and $\mathrm{Var}(R) = \alpha\theta^2$. When reproductive output is standardised to mean 1, $\alpha = 1/I$ and $\theta = I$. Table S5 summarises empirical estimates from strongly polygynous male mammals.

Simulation parameterisation. The high-skew treatment used in the simulations ($\alpha = 0.245$, $\theta = 4.085$; mean ≈ 1, variance ≈ 4.1) closely matches the empirical range observed in several strongly polygynous mammals. Bighorn sheep, southern and northern elephant seals, greater horseshoe bats and Soay sheep independently imply $\alpha \approx 0.22$–$0.29$ (Table S5). Thus, the chosen distribution represents an empirically grounded high-skew scenario. The Gamma distribution was used to generate continuous relative reproductive weights, with parameters moment-matched to empirical reproductive variance.

**Table S5.** Empirical reproductive skew and corresponding moment-matched Gamma distributions.

| Species | Empirical measure | I | Gamma α | Gamma θ (mean = 1) | Reference |
|---|---|---|---|---|---|
| Bighorn sheep | Annual reproductive success | 4.52 | 0.221 | 4.520 | Coltman et al. 2002 |
| Southern elephant seal | Seasonal fertilisation inequality | 4.25 | 0.235 | 4.254 | Galimberti et al. 2002 |
| Northern elephant seal | Breeding success of mature males | 3.91 | 0.256 | 3.910 | Le Boeuf & Reiter 1988 |
| Greater horseshoe bat | Long-term paternity success | 3.70 | 0.270 | 3.700 | Rossiter et al. 2006 |
| Soay sheep | Lifetime breeding success | 3.46 | 0.289 | 3.460 | Coltman et al. 1999 |

Notes. $I = Var(R)/E(R)^2$. For reproductive output standardised to mean 1, $\alpha = 1/I$ and $\theta = I$. Values are representative empirical estimates or values reproduced from comparative compilations cited in the underlying studies. Seasonal reproductive skew can exceed lifetime skew because successful individuals can change among breeding seasons.

### References


Coltman DW, Smith JA, Bancroft DR, Pilkington J, MacColl ADC, Clutton-Brock TH, Pemberton JM. 1999. Density-dependent variation in lifetime breeding success and natural and sexual selection in Soay rams. American Naturalist 154, 730–746.

Coltman DW, Festa-Bianchet M, Jorgenson JT, Strobeck C. 2002. Age-dependent sexual selection in bighorn rams. Proceedings of the Royal Society B 269, 165–172.

Galimberti F, Fabiani A, Sanvito S. 2002. Measures of breeding inequality: a case study in southern elephant seals. Canadian Journal of Zoology 80, 1240–1249.

Le Boeuf BJ, Reiter J. 1988. Lifetime reproductive success in northern elephant seals. In: Clutton-Brock TH (ed.), Reproductive Success: Studies of Individual Variation in Contrasting Breeding Systems. University of Chicago Press.

Rossiter SJ, Ransome RD, Faulkes CG, Dawson DA, Jones G. 2006. Long-term paternity skew and the opportunity for selection in a mammal with reversed sexual size dimorphism. Molecular Ecology 15, 3035–3043.
